# *Xanthomonas euvesicatoria* effector *Xe*AvrRxo1 triggers a Rxo1-mediated defense response in *Nicotiana benthamiana* and its chaperone Xe4429 functions as an antitoxin, an expression repressor, and an enhancer of *Xe*AvrRxo1 secretion

**DOI:** 10.1101/2023.12.02.569727

**Authors:** Zhibo Wang, Changhe Zhou, Tiffany G. Roach, Qi Li, Kunru Wang, Jiamin Miao, Carlos Toro, Shuchi Wu, Yu Tang, Qian Han, Furong Sun, Daniel G. S. Capelluto, Jianyong Li, Bingyu Zhao

**Author notes:** Current address: Regeneron Pharmaceuticals Inc., Basking Ridge, NJ 07920.

## Abstract

*Xanthomonas euvesicatoria* (*X. euvesicatoria*) is the causal agent of bacterial spot disease that threatens pepper and tomato production around the globe. *X. euvesicatoria* gene *Xe4428* encodes a type III effector (T3E) that shares 89.67% amino acid identity with *Xanthomonas oryzae* pv. *oryzicola* (*Xoc*) T3E AvrRxo1. Deletion of *Xe4428* in the genome of *X. euvesicatoria* (strain Xcv85-10) compromised its virulence to infect pepper and *Nicotiana benthamiana* plants. Transient co-expression of Xe4428 and *Rxo1* on pepper and *N. benthamiana* plant leaves results in a robust hypersensitive reaction. Thus, Xe4428, renamed as *Xe*AvrRxo1, is a bona fide orthologue of *Xoc*AvrRxo1 that possesses both virulence and avirulence functions. Expression of *Xe*AvrRxo1 in *E. coli* and *X. euvesicatoria* is toxic to both bacterial cells. Another *X. euvesicatoria* gene *Xe4429*, encodes a putative chaperone of *Xe*AvrRxo1, which can interact with XeAvrRxo1 to suppress its toxicity in *X. euvesicatoria* and *E. coli* bacterial cells. Xe4429 also binds to the promoter region of *XeavrRxo1* and represses its transcription/translation in *X. euvesicatoria* bacterial cells. In addition, expression of Xe4429 can enhance the secretion and translocation of *Xe*AvrRxo1 into plant cells. Therefore, Xe4429 functions as an antitoxin, a transcription repressor, and a type III chaperone that is capable of enhancing the secretion and translocation of *Xe*AvrRxo1 during pathogenesis.

## Introduction

Gram-negative phytobacteria evolved a type III secretion system that can deliver more than 50 type III effectors (T3Es) into plant cells (Jiménez-Guerrero *et al*., 2020). Once inside host cells, T3Es can suppress host immunity and promote bacterial growth or fitness during pathogenesis (Büttner, 2016). Some T3Es are only present in particular bacterial species or specific bacterial strains, while other T3Es are conserved in diverse bacterial species. At the genome level, the conserved *T3E* genes are frequently flanked or adjacent to transposon elements, which is an indication of horizontal transfer of conserved *T3E* genes (Alfano *et al*., 2000; Arnold *et al*., 2001).

We previously identified a T3E, *Xoc*AvrRxo1, from *Xanthomonas oryzae* pv. *oryzicola* (*Xoc*) (Zhao *et al*., 2004). In the genome of *Xoc*, *XocavrRxo1* (ORF1) is adjacent to another gene, *XocavrRxo1*-ORF2, which encodes an atypical molecular chaperone of AvrRxo1 (Han *et al*., 2015). We renamed *Xoc*AvrRxo1-ORF2 as ARC1 (<u>A</u>vrRxo1 <u>R</u>equired <u>C</u>haperone <u>1</u>) (Triplett *et al*., 2016). The avrRxo1 locus is flanked by transposon elements, indicating that it was acquired by horizontal transfer. Indeed, AvrRxo1-homologues have been identified from diverse bacterial species, including a soil myxobacterium *Cystobacter fuscus*, which suggests *Xoc*AvrRxo1 and its homologs may have significant roles even beyond host-pathogen interaction (Triplett *et al*., 2016).

*Xoc*AvrRxo1 triggers *Rxo1*-mediated disease resistance in transgenic rice plants (Zhao *et al*., 2005). *Xoc*AvrRxo1 encodes a NAD kinase that can phosphorylate NAD to form 3’-NADP in both bacterial and plant cells (Shidore *et al*., 2017). Overexpression of *Xoc*AvrRxo1 in yeast, *Escherichia coli,* and *Nicotiana benthamiana* plant cells is toxic, where the toxicity can be blocked by the co-expression of ARC1 (Han *et al*., 2015; Triplett *et al*., 2016). Therefore, ARC1 functions as an antitoxin to suppress the toxicity of *Xoc*AvrRxo1 (Han *et al*., 2015; Triplett *et al*., 2016). A point mutation, D193A, in the substrate-binding site of *Xoc*AvrRxo1, abolishes its toxicity to bacterial cells, and also compromises its ability to trigger the Rxo1-mediated disease resistance in plants (Shidore *et al*., 2017). Another point mutation, T167N, in the ATP-binding motif, also suppresses the toxicity of AvrRxo1 but only partially abolishes its NAD kinase enzyme activity and virulence function (Shidore *et al*., 2017). Other than the role of antitoxin, the biological and biochemical functions of ARC1 are not characterized. For unknown reasons, deletion of either *XocavrRxo1* alone or *avrRxo1*-*ARC1* together was not successful, which limits us to further investigate the biological function of AvrRxo1-ARC1 in *Xoc*. Therefore, we attempted to characterize AvrRxo1 and ARC1 homologs in another bacterial pathogen, *Xanthomonas euvesicatoria*.

*X. euvesicatoria* is the causal pathogen of bacterial spot (BS) disease of pepper and tomato. The BS disease is one of the most economically important pepper and tomato diseases prevalent in regions with warm and high humidity climates (Potnis *et al*., 2015). The causal pathogen was originally named *Xanthomonas campestris vesicatoria* (*Xcv*). Biochemical characterization, genome sequencing, and phylogenic analysis of xanthomonads, isolated from pepper and tomato plants, led to reclassifying of the *Xcv* strains into four subgroups: *X. euvesicatoria* (A), *X. vesicatoria* (B), *X. perforans* (C), and *X. gardneri* (D) (Schwartz *et al*., 2015; Stall *et al*., 1994; Vauterin *et al*., 1995). Most *Xanthomonas* strains infecting pepper belong to group A, which can be further classified into 11 races of *X. euvesicatoria* based on their pathogenesis in pepper plants carrying different disease resistance genes (Jones *et al*., 2004; Potnis *et al*., 2015; Stall *et al*., 2009). Thus far, six BS resistance genes have been identified, most of which are race-specific (Vallejos *et al*., 2010).

One *X. euvesicatoria* strain, Xcv85-10, originally isolated in Florida, U.S., can infect both pepper and tomato plants. Xcv85-10 has been selected as a model strain for studying the pepper/*X. euvesicatoria* interactions (Thieme *et al*., 2005). Analysis of the genome sequence of Xcv85-10 revealed >20 putative T3Es (Thieme *et al*., 2005). Several T3Es in strain Xcv85-10 have been functionally characterized (Popov *et al*., 2016; Roden *et al*., 2004; Teper *et al*., 2016; Zhao *et al*., 2011). However, the biological functions of most T3Es in Xcv85-10 are still unclear. One predicted T3E Xe4428 in Xcv85-10 is identified as the *Xoc*AvrRxo1 homolog (Triplett *et al*., 2016; Zhao *et al*., 2004). Another *X. euvesicatoria* gene *Xe4429*, located downstream of *Xe4428*, shares high sequence homology with *ARC1*. Thus far, the biological and biochemical functions of Xe4428 and Xe4429 have not yet been characterized.

In this study, we demonstrated that Xe4428 is a functional orthologue of *Xoc*AvrRxo1, which can trigger the *Rxo1*-mediated defense responses in *N. benthamiana*. Therefore, we renamed Xe4428 as *Xe*AvrRxo1. *Xe*AvrRxo1 also has a significant virulence function that can promote ***X. euvesicatoria*** proliferation on both pepper and *N. benthamiana* plants. We further revealed that *Xe4429* encodes an antitoxin that can suppress the toxicity of *Xe*AvrRxo1. In addition, we demonstrated that Xe4429 binds to the promoter region of *Xe*AvrRxo1 and suppresses its expression in *X. euvesicatoria* bacterial cells. An adenylate cyclase reporter assay revealed that the expression of Xe4429 could enhance the secretion and translocation of *Xe*AvrRxo1. Taken together, Xe4429 functions as an antitoxin, a transcription repressor, and a putative chaperone that is capable of prompting the secretion of a specific T3E in *X. euvesicatoria*.

## Methods and Materials

### Plant materials and growth conditions

*N. benthamiana*, pepper (*Capsicum annuum*, cv. Early Calwonder, (ECW)) seeds were germinated and grown in Sunshine® Mix #1 in a growth chamber (14h/10h light/dark cycle at 25°C/20°C). The 6-week-old tobacco, pepper, and tomato plants were used for the experiments detailed herein.

### Bacterial growth

*E. coli* strains *DH5α* and *C41(DE3)* (Lucigen, Middleton, WI) were grown on Luria agar medium at 37 ℃. *Agrobacterium tumefaciens GV2260* strain and *X. euvesicatoria* were grown on Luria agar medium and NYG medium at 28 ℃, respectively (Zhao *et al*., 2011). *E. coli* antibiotic selections used in this study were as follows: 50 μg/ml kanamycin, 100 μg/ml carbenicillin, 100 μg/ml spectinomycin, 34 μg/ml chloramphenicol, and 50 μg/ml gentamycin. *X. euvesicatoria* and *A. tumefaciens* antibiotic selection were 100 μg/ml rifampicin and/or 50 μg/ml kanamycin.

### Gene cloning, site-directed mutagenesis, plasmid construction, and Agrobacterium-mediated transient assay

The open reading frames (ORFs) of *XeavrRxo1* and *Xe4429*, *XeavrRxo1*-*Xe4429*, native promoter-*XeavrRxo1*-*Xe4429*, and the native *XeavrRxo1* promoter only, were amplified from the genomic DNA of Xcv85-10. The *Rxo1* ORF was amplified from the genomic DNA of maize cv. B73. All PCR primers with annotations are listed in Table S1. The ORF of NanoLuc® Luciferase was amplified from pNanoLuc (Promega) and cloned into pDonr207 (Thermo Fisher Scientific). Other PCR products were cloned into pENTR/D-TOPO (Thermo Fisher Scientific). pENTR/D-*XeavrRxo1*(D222T), pENTR/D-native promoter-*XeavrRxo1*(D222T)-*Xe4429*, pENTR/D-native promoter-*XeavrRxo1*(D222T)-*Xe4429*, pENTR/D-*Rxo1*(D291E), were generated *via* site-direct-mutagenesis using the primers listed in Table S1 (Han *et al*., 2015). The genes/fragments in donor vectors were subcloned into the pEarleyGate101 using a LR^®^ Gateway cloning kit (Thermo Fisher Scientific) (Earley *et al*., 2006). The plant expression constructs were transformed into *A*. *tumefaciens* strain *GV2260*. *Agrobacterium* strains harboring different constructs were infiltrated into leaf mesophyll tissues for transient protein expression (Traore *et al*., 2019). The transient assay pictures were taken at 48 hours post-inoculation. Plant-expressed proteins were detected by Western blot analysis following the procedure as previously described (Liu *et al*., 2020). The expressed proteins were detected by using anti-HA-HRP (Roche) at a dilution of 1:5,000. The ion leakage was analyzed as previously described (Liu *et al*., 2015). For each treatment, at least three leaf disks (1 cm^2^) and three replicates were collected in ddH_2_O. The solution conductivity (C1) was measured with a conductivity meter (SR60IC, VWR, Radnor, PA, USA). The leaf samples were autoclaved, and the conductivity of the solution containing the killed tissue was measured (C2). The relative leaf electrolyte leakage was calculated using the formula EL (%) = (C1/C2) ×100.

### RNA isolation and real-time PCR

Total RNA was extracted from *N. benthamiana* leaves using TRIzol reagent (Thermo Fisher Scientific) according to the manufacturer’s instructions. Any DNA residue was eliminated by treating with UltraPure DNase I (Thermo Fisher Scientific). The integrity and quantity of total RNA were determined by electrophoresis in 1% agarose gel, and a NanoDrop ND-1000 spectrophotometer (NanoDrop Technologies, Wilmington, DE). cDNA synthesis was performed using the SuperScript III First-Strand RT-PCR Kit (Thermo Fisher Scientific) with an oligo-dT primer based on the manufacturer’s instructions. Real-time PCR was conducted with cDNA 20 using the Quantitect SYBR Green PCR kit (Qiagen) according to the manufacturer’s protocol. Oligo primers are listed in Table S1. The tobacco actin gene was used as a reference gene, and data is presented as ΔΔCT.

### Marker-exchange mutagenesis and the complementary constructs

To delete the *XopQ* gene from the genome of Xcv85-10, we amplified the *XopQ* upstream and downstream DNA fragments using primers listed in Table S1. DNA fragments were cloned into PCR8-Km that carries a *kanamycin* (Km) resistance gene cassette (Liu *et al*., 2016), resulting in PCR8-up-Km-down. This construct was used for subcloning the up-Km-down fragment into a Gateway-compatible vector pLVC18L-SacB/R-Des (Liu *et al*., 2016; Zhao *et al*., 2011). The derived construct pLVC18L-KO-XopQ was used for marker-exchange mutagenesis in Xcv85-10 (Zhao *et al*., 2011). The Km resistance gene was removed by using a Flipase expression vector (Levy *et al*., 2018). This marker-free *X. euvesicatoria* mutant was designated as *Xe*ΔXopQ (Figure S1). A similar procedure was used to delete the *XeavrRxo1-*Xe4429 locus in Xcv85-10 or *Xe*ΔXopQ (Figure S1). The derived marker-free mutants were designated *Xe*Δ*XeavrRxo1* and *Xe*Δ*XopQ*Δ*XeavrRxo1*, respectively.

A broad host-spectrum vector pBMTBX-2 (Prior *et al*., 2010) was converted into a destination vector by cloning the *ccdB* cassette (Thermo Fisher Scientific Inc.) into the *Nco*I site. The derived vector was designated as pBMTBX-2-Des. The ORFs of *XeavrRxo1*, *XeavrRxo1*(D222T), *XeavrRxo1*-*Xe4429*, and *Xe4429* were subcloned into pBMTBX-2-Des via LR^®^ cloning. The donor vector carrying native promoter-*XeavrRxo1*-*Xe4429* or native promoter-*XeavrRxo1*(D222T)-*Xe4429* was subcloned into another broad-host spectrum vector pVSP61-Des (Traore *et al*., 2019) via LR^®^ cloning. The derived constructs were named as pNative pro-*XeavrRxo1*-*Xe4429* and pNative pro-*XeavrRxo1*(D222T)-*Xe4429*, and were used to complement the *XeavrRxo1* deletion in *Xe*Δ*XeavrRxo1* or *Xe*Δ*XopQ*Δ*XeavrRxo1*.

A DNA fragment was amplified with primers “Xe4426-Sac2 For” and “Xe4428_353Sac2 Rev” from the genome DNA of Xcv85-10, and cloned into the *Sac*II site of pENTR-native promoter-*XeavrRxo1*(D222T). The DNA fragment was subcloned into a Gateway-compatible vector pLVC18L: *Cya*-Des via LR cloning (Zhao *et al*., 2011). The derived construct was designated as pLVC18L-D222T: *Cya* that was integrated into the genome of *Xe*Δ*XopQ*Δ*XeavrRxo1* via marker-exchange mutagenesis.

### Bacterial growth curve assay

Bacterial proliferations on inoculated *N. benthamiana* and *C. annuum* plants were assessed via standard growth curve assays (Zhao *et al*., 2011). In brief, bacterial strains were cultivated on NYG agar plates at 28°C for two days. Bacterial cells were collected and suspended in 10 mM MgCl_2_ with 0.02% Silwet L77. For spray inoculation, a bacterial inoculum was diluted to 2 x10^8^ CFU ml^-1^ and sprayed on plant leaves using a spray bottle. Inoculated plants were maintained under 95% relative humidity overnight and then cultivated under 14 h light/10 h dark cycles for six days. Leaf discs (990 mm^2^) were randomly sampled from inoculated leaves for growth curve assays. The sampled leaf discs were ground in 10 mM MgCl_2_ and plated on NYG medium supplemented with appropriate antibiotics. The bacterial colony forming units (CFU) were counted, and the bacteria proliferation ratio was calculated as Log_10_ CFU/cm^2^. All growth curve assays underwent three biological repeats with at least three technical replicates.

### Monitoring the growth rates of E. coli and X. euvesicatoria strains expressing XeavrRxo1

The pBMTBX-*XeavrRxo1*, pBMTBX-*XeavrRxo1*(D222T), pBMTBX-*XeavrRxo1*-*Xe4429*, and pBMTBX-*NanoLuc* (negative control) were transformed into *E. coli* strain *C41(DE3)*. Bacterial strains were grown at 37°C in LB liquid medium supplemented with 50 μg/ml Kanamycin. The bacterial culture was diluted to OD_600_ = 0.1 as the starting point, and gene expression was induced by adding 0.4 % arabinose. Bacterial cultures, supplemented with 0.4 % glucose, were used as non-induction controls. The same set of constructs was transformed into *Xe*Δ*XeavrRxo1* for growth rate assay. In addition, pNative pro-*XeavrRxo1*-*Xe4429* and pNative pro-*XeavrRxo1*(D222T)-*Xe4429*, were also transformed into *Xe*Δ*XeavrRxo1* for monitoring the growth rates. The *X. euvesicatoria* strains were grown in liquid NYG medium, and the bacterial growth was monitored using a spectrophotometer (Beckman Coulter, Model DU 800).

### Cloning, expression, and purification of XeAvrRxo1 and Xe4429

The *XeAvrRxo1*-*Xe4429* and *Xe4429* genes cloned in pTopoENTR were subcloned into a Gateway-compatible pGEX4T-1 destination vector via LR^®^ cloning (Han *et al*., 2015). The plasmids were transformed into *E. coli C41* cells (Lucigen). XeAvrRxo1:Xe4429 and Xe4429 proteins were expressed and purified following a procedure as previously described (Han *et al*., 2015). Protein purity was evaluated by SDS–PAGE. The protein concentration was determined by a protein assay kit (Bio-Rad) using bovine serum albumin as the standard (Han *et al*., 2015).

### Electrophoretic mobility shift assay

Electrophoretic mobility shift assay (EMSA) was performed as previously reported (Van Eck *et al*., 2014). The promoter DNAs were amplified from pENTR-*XeavrRxo1* promoter using either the FAM (645)-labeled M13For and M13Rev or non-labelled M13For and M13Rev primers (Integrated DNA technologies Inc. Iowa). DNA concentrations were determined using a ND-1000 spectrophotometer (NanoDrop Technologies). The FAM-labeled DNA fragment of *XeavrRxo1* promoter was mixed with either *Xe*AvrRxo1:Xe4429 or Xe4429 proteins, along with different amounts of the unlabeled competitor DNA fragment in a reaction buffer (10 mM Tris-HCl, 100 mM KCl, 1 mM EDTA, 0.1 mM DTT, 5 % glycerol, 0.01 mg/mL BSA, pH 7.5) for 20 min at room temperature. Reactions were separated on 1 % agarose gel in 1xTAE buffer. The fluorescence signal was captured using a gel scanner, Typhoon FLA7000, equipped with a 635 nm laser filter (GE Healthcare, Piscataway, NJ).

### Isothermal titration calorimetry interactions

Isothermal titration calorimetry (ITC) titrations were performed using a Malvern Panalytical Microcal PEAQ instrument. The ITC buffer consisted of 20 mM HEPES (pH 7.3) and 150 mM KCl. To generate an 81-bp *XeavrRxo1* promoter DNA fragment, two complementary oligos (Xe4428pro-81For/Rev, Table S1) were synthesized (Integrated DNA Technologies Inc). The oligos were annealed, ethanol-precipitated, and the DNA pellet was redissolved in ITC buffer. The *XeavrRxo1* promoter DNA was titrated into Xe4429 at concentrations of 1 mM and 35-50 µM, respectively. Concentrations were determined using a Thermo Scientific NanoDrop One instrument. The dsDNA program was used for DNA concentration analysis, ensuring intact double strands with a 260:280 nm ratio of ∼1.8. Protein concentration was determined based on absorbance at 280 nm, with baseline correction using a reading at 340 nm. Prior to each experiment, the ITC instrument was equilibrated to 25°C, and the deferential power was set to 10 µcal/sec. Twelve injections of 3 µL each were used, with an initial test injection of 0.4 µL to ensure system stability. Malvern’s MicroCal PEAQ analysis software was used for analyzing the binding traces, considering a one-site binding model and subtracting the heats of dilution from *XeavrRxo1* promoter DNA titrated into buffer. The dissociation and Gibbs free energy constants are reported as the average of three independent experiments.

### Expressional regulation of XeavrRxo1 mediated by Xe4429

The transcriptional regulation activity of Xe4429 was detected using a *XeavrRxo1 promoter* (*XeavrRxo1*pro)-*NanoLuc* reporter construct as described below. The *XeavrRxo1*pro-*NanoLuc* fragment was amplified by using an overlap PCR approach (Table S1). The purified PCR product was cloned into the *Xba*I site of pBMTBX-2-Des to generate pBMTBX-*XeavrRxo1*pro-*NanoLuc*. The *Xe4429*, *XeAvrRxo1*-*Xe4429*, or *GFP* genes, previously cloned in donor vectors, were subcloned into pBMTBX-*XeavrRxo1*pro-Nalouc via LR^®^ cloning. The *XeavrRxo1*pro-*NanoLuc* gene was cloned into pVSP61-Des (Traore *et al*., 2019) as a control. The pBMTBX-*NanoLuc* was also used as a control. The key elements of these constructs are illustrated in Figure 7A. The reporter constructs were transformed into *XeρavrRxo1* and used for *NanoLuc* reporter assay *in vitro*. In brief, bacteria were cultured on the NYG liquid medium supplemented with kanamycin (50 µg/ml). The overnight culture was collected and adjusted to OD_600_ = 0.4 with 10 mM MgCl_2_. The bacterial cells carrying pBMTBX-*NanoLuc* were further diluted (1:500) because the signal was too strong. Gene expression was induced with 0.4 % arabinose for 6 hours, and 0.4 % glucose was used as a control. The NanoLuc substrate (1:1,000 dilution) (Promega) was added to the bacterial cultures. After incubation for 10 min, the bioluminescence signals were detected using a CCD camera of a Gel Doc^TM^ XR+ System (Bio-Rad).

The bacterial strains harboring the same set of constructs as described above were also used for inoculating the pepper leaves via infiltration. The NanoLuc substrate was applied to the leaves 24 hours post-inoculation. The bioluminescence signals in pepper leaves were detected using a Gel Doc^TM^ XR+ System.

### cAMP assay from pepper leaf tissue inoculated with X. euvesicatoria strains

*X. euvesicatoria* cells were adjusted to 4 x 10^8^ CFU ml^-1^ with 10 mM MgCl_2_ supplemented with either 0.4 % glucose or 0.4 % arabinose. The bacterial inoculum was infiltrated into pepper leaf tissue with a blunt-end syringe. Leaf disks (1 cm^2^) were collected at 8 h post-infiltrations with *X. euvesicatoria* strains harboring the indicated *Cya* reporter genes. Leaf disks were collected in three replicates for each treatment. cAMP extractions and protein measurements were performed as previously described (Casper-Lindley *et al*., 2002; Zhao *et al*., 2011). cAMP was measured by using a cAMP enzyme immunoassay kit (AAT Bioquest, Inc.) according to the manufacturer’s instructions.

### CD spectroscopy

Far-UV CD spectra were collected using purified Xe4429 (10 μM) in 5 mM Tris-phosphate (pH 6.8) and 100 mM potassium fluoride on a Jasco J-815 spectropolarimeter equipped with a PFD-425 S temperature-control unit. Five accumulated spectra were recorded in a 0.1-cm quartz cuvette at 25°C with 1-sec-response as well as 0.5-nm-data pitch at a scan speed of 50 nm/min. Buffer backgrounds were used to subtract protein spectra.

### Homology modeling and structural comparison of Xe4429 with WhiA and ARC1

A 3D model of Xe4429 was built using Swiss model (https://swissmodel.expasy.org/) and optimized by potential energy minimization using Gromacs 5.1.4 package. The quality of the model was analyzed using Molprobity Ramachandran Analysis (Williams *et al*., 2018). The 3D alignment of the structures of Xe4429, *Xoc*AvrRxo1-ORF2 (ARC1), and WhiA was done using STRAP (Gille and Frömmel, 2001) and visualized by Pymol (http://www.pymol.org).

### Statistical data analysis

Analytical experiments were performed a minimum of three times with at least three technical replicates. Statistical significance was based on a t-test for paired comparisons or the Tukey–Kramer honestly significant difference test for multiple comparisons. Data was analyzed using JMP Pro14. Values of P<0.05 were considered significant.

## Results

### Xe4428 is a homolog of XocAvrRxo1 that can trigger the Rxo1-mediated defense response in N. benthamiana

We previously determined the three-dimensional structure of *Xoc*AvrRxo1 along with its putative chaperone ORF2 (ARC1) (Han *et al*., 2015). Xe4428 shares 89.67% amino acid sequence identity with *Xoc*AvrRxo1 (Figure 1A). Here, we built a 3D model of Xe4428 using *Xoc*AvrRxo1 (pdb code:4z8q) as a template. The optimized model has GMQE and QMEAN scores of 0.57 and 0.31, respectively. Molprobity Ramachandran Analysis shows that 98.18% of all residues were in favored regions, and 0.31% of residues were in outliers. The 3D alignment of Xe4428 and *Xoc*AvrRxo1 is shown in Figure 1B. Therefore, Xe4428 shares significant structure homology with *Xoc*AvrRxo1, and consequently, we renamed it as *Xe*AvrRxo1.

**Figure 1.**
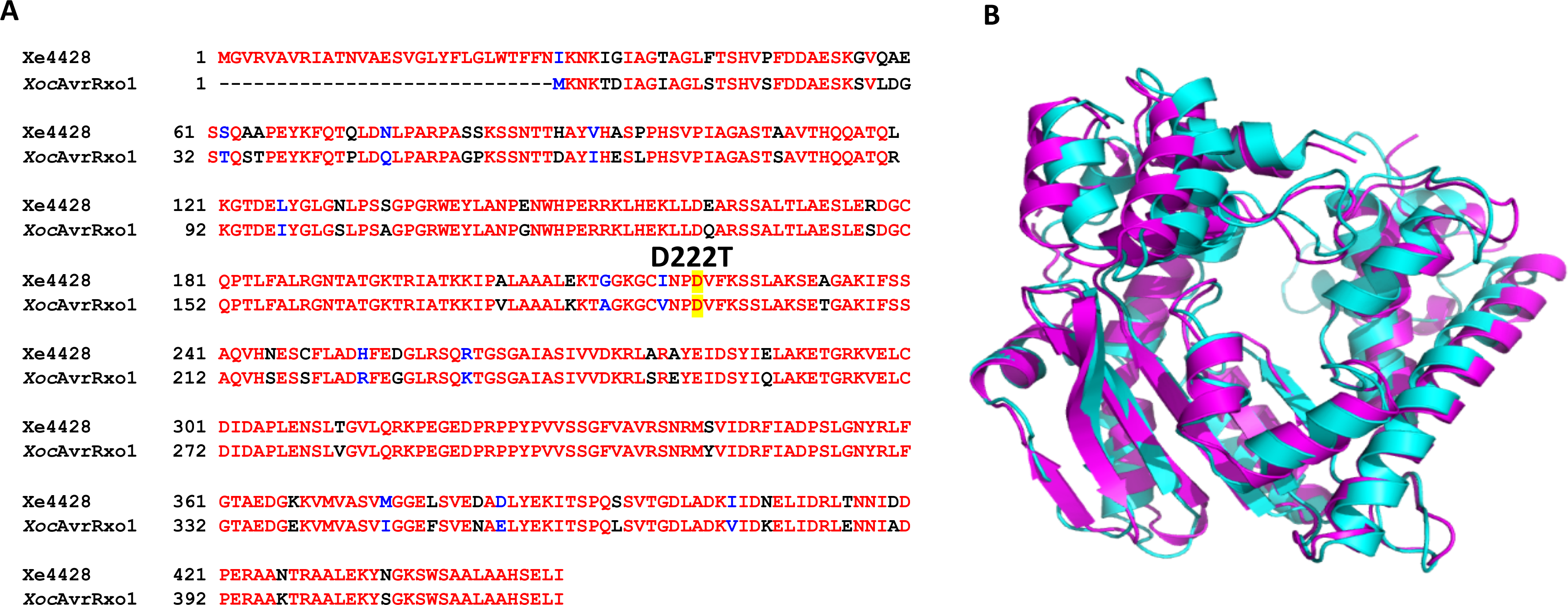
The predicted three-dimensional structure of Xe4428. (A). Xe4428 shares 84% amino acid sequence identity with *Xoc*AvrRxo1. The conserved amino acids are in red; the polymorphic amino acids are in blue (semi-conserved) or black (different). A conserved aspartic acid residue that is required for NAD enzymatic activity is highlighted, and the corresponding mutation, D222T, is indicated. (B). The predicted three-dimensional structure of Xe4428 (purple) was overlaid with that corresponding to *Xoc*AvrRxo1 (4z8q, blue).

To test whether *Xe*AvrRxo1 can trigger an Rxo1-mediated defense response in *N. benthamiana* plants, we performed *Agrobacterium*-mediated transient expression of *Rxo1* and *XeavrRxo1* on *N. benthamiana* plant leaves. Wild type *Xe*AvrRxo1, but not a substrate-binding mutant *Xe*AvrRxo1 (D222T), triggered a strong hypersensitive response on transformed plant leaves (Figure 2A). As controls, a mutant of Rxo1, Rxo1(D291E), could not recognize either wild type or mutant *Xe*AvrRxo1 to trigger cell death in *N. benthamiana* (Figure 2A). Western blotting confirmed both wild-type and mutant proteins were equally expressed in transformed plant cells (Figure 2B). The strong cell death promoted by the Rxo1/*Xe*AvrRxo1 also resulted in an increased ion leakage (Figure 2C). In addition, the expression of defense-related genes *NbPR5* and *NbPR10,* but not *NbPR1*, were dramatically induced by the interaction of Rxo1 and *Xe*AvrRxo1 (Figure 2D). Interestingly, another defense-related gene, *NbPR4*, encodes a putative chitinase and is significantly downregulated by the interaction of Rxo1/*Xe*AvrRxo1 (Figure 2D). Surprisingly, co-expression of *Rxo1*(D291E) with *XeavrRxo1* could induce *Nb*PR4 (Figure 2D). Nevertheless, we confirmed that *Xe*AvrRxo1 could trigger the Rxo1-mediated defense responses in *N. benthamiana*.

**Figure 2.**
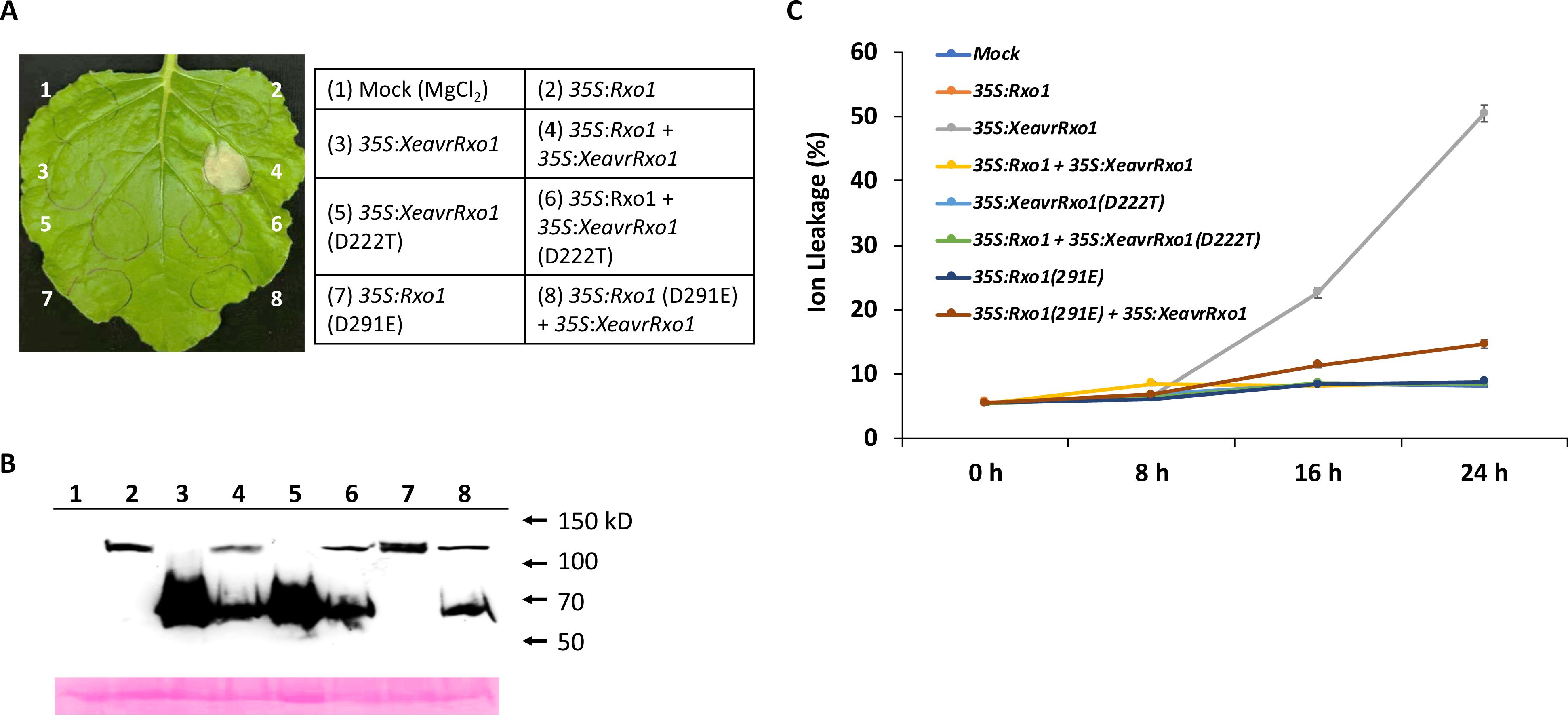
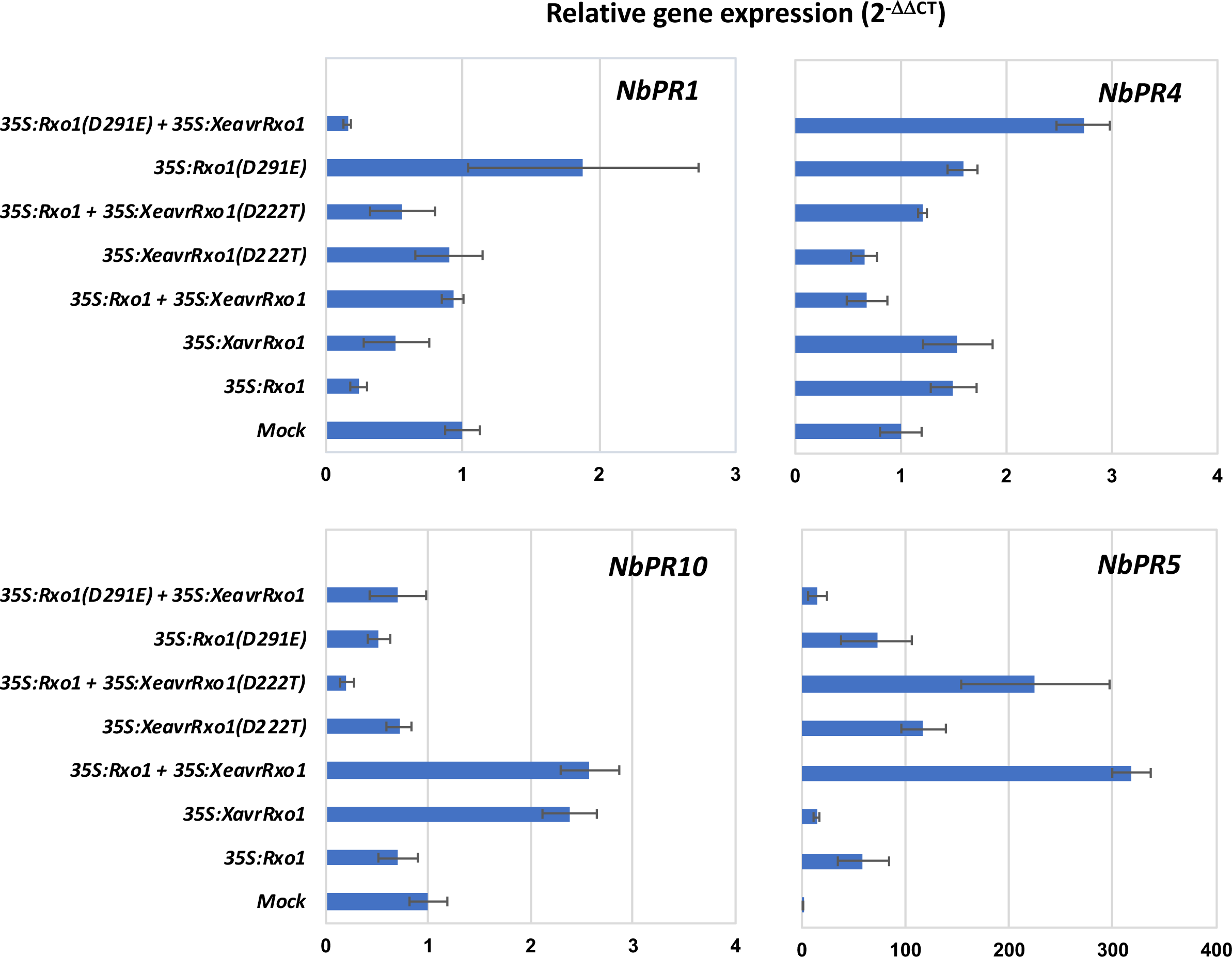
Transient expression of *XeavrRxo1* triggers the *Rxo1*-mediated defense response in *N. benthamiana*. (A) Wild type, but not mutant *XeavrRxo1,* triggered *Rxo1*-mediated cell death in *N. benthamiana*. Bacterial concentrations were adjusted to 4 x 10^8^ CFU ml^-1^ (single inoculation) or 2 x 10^8^ CFU ml^-1^ (co-inoculation). Picture was taken two days post-inoculation. (B) The targeting proteins fused with HA epitope tag as listed in (A) were detected with anti-HA. The blot membrane was stained with Ponceau S to confirm equivalent loadings. (C) Ion leakage of leaf samples, as indicated in (A), was monitored at 0, 8, 16, and 24 h post-inoculation. (D) The expressions of defense-related genes *NbPR1*, *NbPR4*, *NbPR5,* and *NbPR10* were monitored by qPCR. Leaf samples were collected at 24 h post *Agrobacterium*-mediated transient assays as indicated in (A). Experiments were repeated three times with similar results, and data are shown as mean ± s.e.

### XeAvrRxo1 possesses a virulence function that can enhance X. euvesicatoria proliferation in both pepper and tobacco plants

A recent report suggests that deleting the T3E gene *XopQ* in *X. euvesicatoria* alters its host specificity and allows the mutant *X. euvesicatoria* strain to infect *N. benthamiana* plants (Schwartz *et al*., 2015). To characterize the virulence function of *Xe*AvrRxo1 on *N. benthamiana*, we generated isogenic *X. euvesicatoria* mutant strains that carry a deletion of *XopQ* alone or a double-deletion of both *XeXopQ* and *XeavrRxo1-Xe4429*.

The isogenic *X. euvesicatoria* strains (*XeΔXopQ* and *XeΔXopQΔ*(*XeavrRxo1-Xe4429*) were able to infect *N. benthamiana* and pepper plants. From hereon, the mutant *Xe* strain with deletion of *XeavrRxo1-Xe4429* is referred to as *XeΔXeavrRxo1* for simplicity. The *Xe*Δ*XopQ*Δ*XeavrRxo1* strain, when inoculated, exhibited markedly lower population levels than XeΔXopQ on both *N. benthamiana* and pepper plants (Figures 3 A and B). This result suggests that *Xe*avrRxo1 has a virulence function that can promote bacterial proliferation in inoculated plants. The *XeΔXopQΔXeavrRxo1* mutant strain was transformed with a plasmid carrying either wild type *XeavrRxo1-Xe4429* or mutant *XeavrRox1*(D222T)-*Xe4429* genes, where the expressions of both genes are driven by their native promoters. A growth curve assay revealed that *X. euvesicatoria* strain with wild type, but not D222T of *XeavrRxo1,* was able to grow to a population level similar to *XeΔXopQ* on both *N. benthamiana* and pepper plants (Figures 3 A and B). Therefore, *Xe*AvrRxo1 possesses a strong virulence function that may require its NAD kinase enzymatic activity.

**Figure 3.**
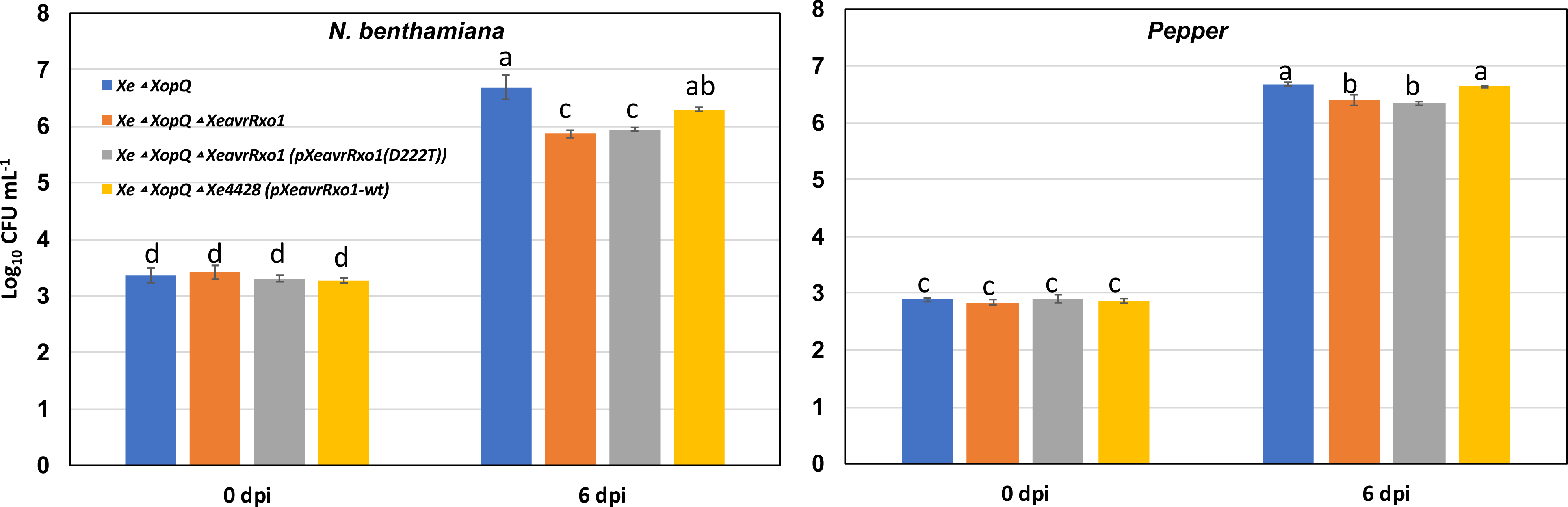
*XeavrRxo1* possesses a virulence function that can enhance *X. euvesicatoria* proliferation on pepper and *N. benthamiana* plants. *In planta,* bacterial growth of *XeρXopQ*, *XeρXopQρXeavrRxo1*, *XeρXopQρ XeavrRxo1* (pNative pro-*XeavrRxo1*-*Xe4429*), and *XeρXopQρ XeavrRxo1* (pNative pro-*XeavrRxo1*(D222T)-*Xe4429*) was measured at 0 and 6 days after infiltration in leaves of *N. benthamiana* (A) and pepper (B). The starting inoculum of *Xe* strains was adjusted to 1 × 10^5^ CFU ml^−1^. Experiments were repeated three times with similar results and data are shown as mean ± s.e. WT: wild type gene. Different letters indicate significant statistical differences (*p*-value < 0.05).

### Expression of XeavrRxo1 is toxic to X. euvesicatoria and E. coli, and the toxicity can be suppressed by co-expression with Xe4429

Our previous results demonstrated that the expression of *Xoc*AvrRxo1 in *E. coli* is toxic, and toxicity can be suppressed by co-expression with ARC1(Triplett *et al*., 2016). To test whether *Xe*AvrRxo1 is also toxic to bacterial cells and whether Xe4429 can block the toxicity of *Xe*AvrRxo1, we employed an arabinose-inducible expression system to express wild-type *XeavrRxo1* or D222T with or without the expression of Xe4429 in *E. coli* and *X. euvesicatoria* cells. The expression of *XeavrRxo1* but not D222T, or co-expression of *XeavrRxo1* with *Xe4429* inhibits *E. coli* growth (Figure 4A). As a control, when bacterial cells were grown in LB medium without arabinose, all bacterial strains, except the strain carrying *XeavrRxo1* alone, displayed a similar growth rate. It is possible that the leaking expression of *XeavrRxo1* might reduce bacterial growth (Figure 4A). Nevertheless, we conclude that expression of *Xe*AvrRxo1 is toxic to *E. coli*, and that co-expression with Xe4429 could reduce its toxicity. The point mutation, D222T, located in the substrate-binding site, also abolished the toxicity of *Xe*AvrRxo1. The same set of expression constructs was also transformed into a mutant *X*.

**Figure 4.**
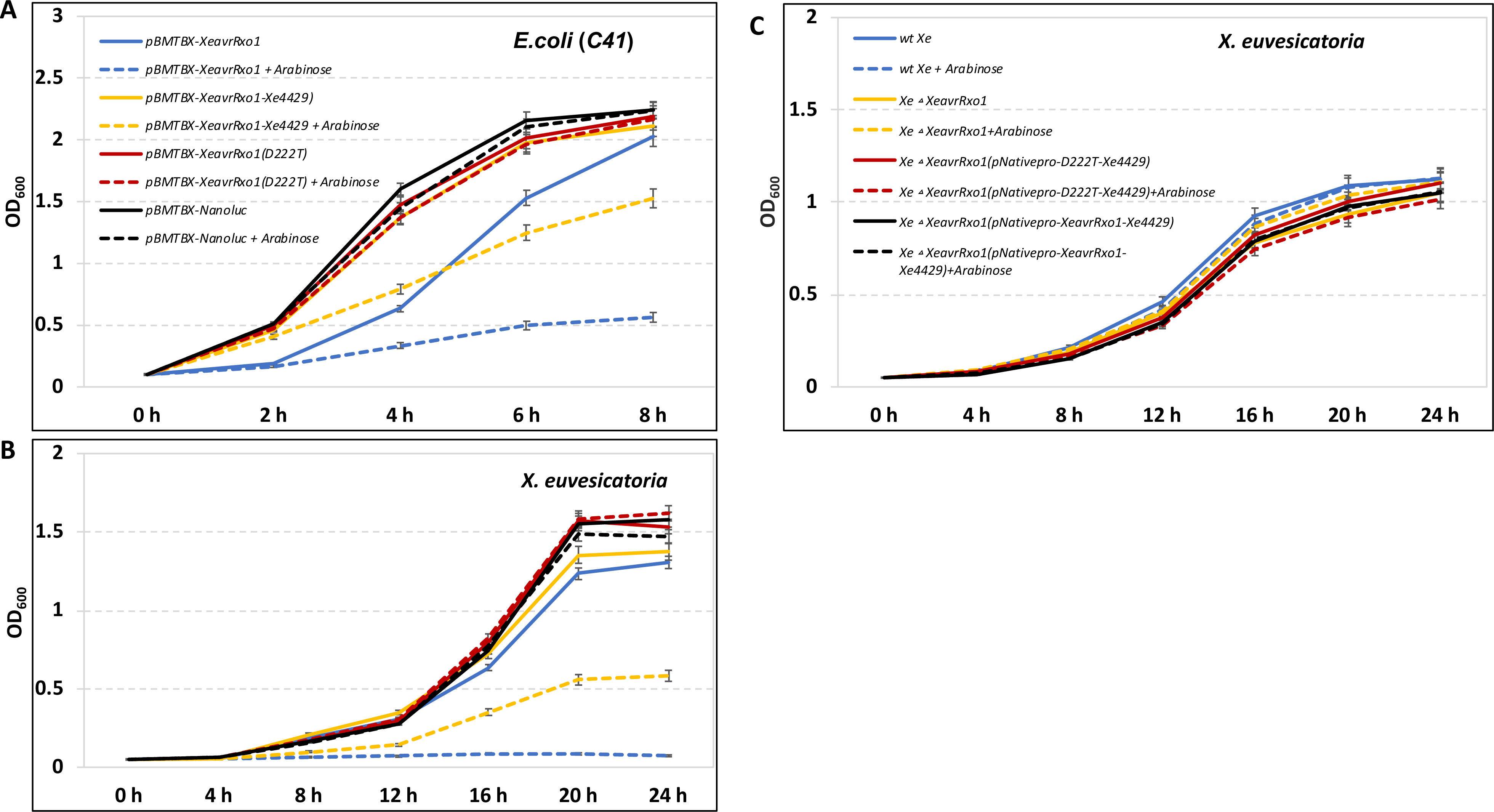
Expression of *XeAvrRxo1* is toxic to *X. euvesicatoria* and *E. coli*, and its toxicity can be suppressed by *Xe4429*. The growth rates of *E. coli* C41 (A) and strain *XeΔavrRxo1* (B) strains. The bacterial strains carry the following expression constructs: (1) pBMTBX-*XeavrRxo1*, (2) pBMTBX-*XeavrRxo1*-*Xe4429*, (3) pBMTBX-*XeavrRxo1*(D222T), and (4) pBMTBX-*NanoLuc*, were grown in liquid medium supplemented with and without arabinose. (C) The growth rates of *Xe* strains, including wild type *Xe* (Xcv85-10), *XeΔavrRxo1*, *XeΔavrRxo1* (pNative pro*-XeavrRxo1*-*Xe4429*), and *XeΔavrRxo1* (pNative pro-*XeavrRxo1*(D222T)-*Xe4429*) grown in liquid NYG medium at 28°C, supplemented with or without arabinose. Experiments were replicated three times and data are shown as mean ± s.e.

The same set of expression constructs was also transformed into a mutant *X*. *euvesicatoria* strain, *XeΔavrRxo1,* where both *XeavrRxo1* and *Xe4429* genes were deleted. Transformed *X*. *euvesicatoria* strains were used for a growth curve assay. Inducible expression of wild-type, but not D222T of *XeavrRxo1*, could strongly inhibit the proliferation of *Xe* bacterial cells. Surprisingly, co-expression with Xe4429 still showed a slower growth rate than that of other controls (Figure 4B). To rule out the possibility that arabinose or the arabinose-inducible expression elements could alter the bacterial growth rates, we also compared the growth rates of *X. euvesicatoria* strains with plasmids carrying native promoter-*XeavrRxo1-*Xe4429 genes. As shown in Figure 4C, wild-type *X. euvesicatoria* strain Xcv85-10, *XeΔavrRxo1*, *XeΔavrRxo1* (pNative pro-*XeavrRxo1-Xe4429*), *XeΔavrRxo1* (pNative pro-*XeavrRxo1*(D222T)*-Xe4429*) strains have almost identical growth rates in the conditions with or without the supplementation of arabinose. Therefore, we conclude that when *XeavrRxo1* and *Xe4429* are co-expressed by the native promoter in *X. euvesicatoria*, the toxicity of *Xe*AvrRxo1 could be completely blocked by Xe4429. Thus, Xe4429 is a functional orthologue of ARC1.

### Xe4429 shares a limited structure homology with a transcription regulator WhiA

To investigate the biochemical functions of Xe4429 further, we predicted its three-dimensional structure through homology-based modeling. Xe4429 shares 83% of its identity with ARC1 (Figure 5). A three-dimensional model of Xe4429 was built using ARC1 (PDB:4z8v) as a template (Han *et al*., 2015) (Figure 5A). The final model has GMQE and QMEAN scores of 0.88 and 0.14, respectively. Molprobity Ramachandran analysis shows that 86.5% of all residues are in favored regions, 100% of all residues are in allowed regions, and no residues are outliers. A purified Xe4429 protein was also analyzed by circular dichroism spectroscopy (Figure S2), which showed that Xe4429 is largely α-helical, in agreement with previous observations (Yang *et al*., 1986).

**Figure 5.**
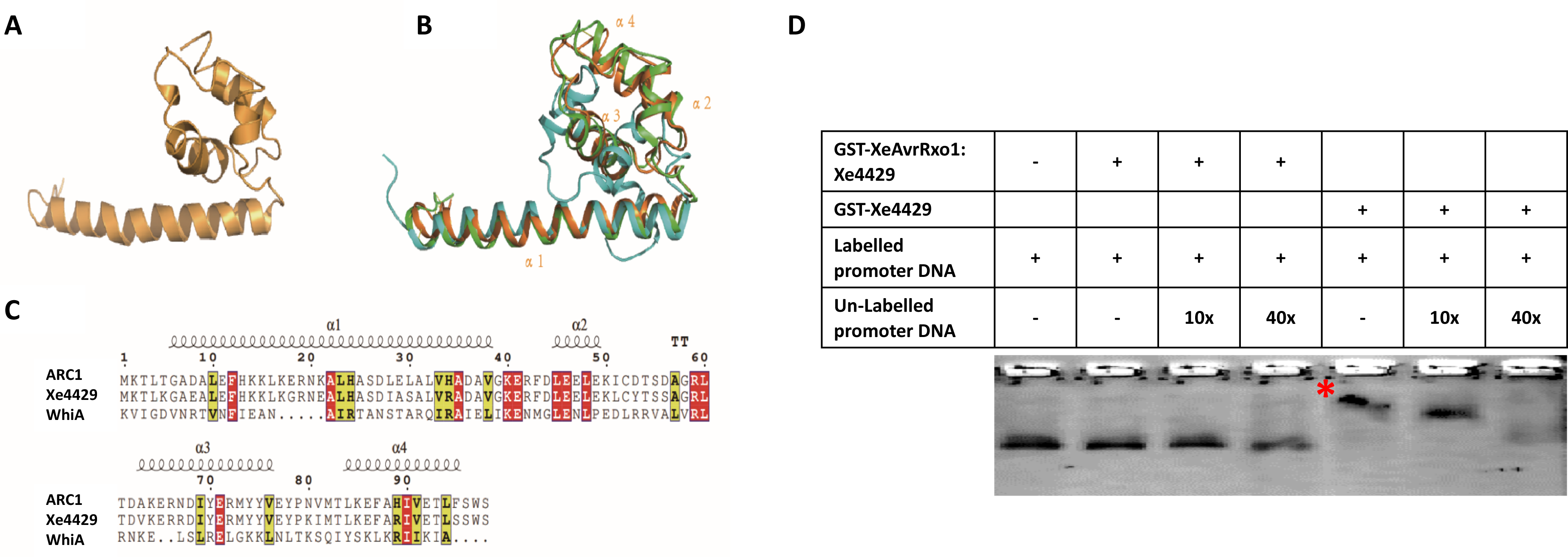
The predicted Xe4429 structure shares homology with a transcription regulator DUF199/WhiA (3hyi-Cter), and purified Xe4429 protein binds to the promoter of *XeavrRxo1*. (A) Optimized homology model of Xe4429 is drawn in cartoon backbone representation. (B) Visualization of the three-dimensional structural alignment of *Xoc*AvrRxo1 ORF2 (ARC1), DUF199/WhiA, and three-dimensional model of Xe4429. Cartoon backbone representation of the three-dimensional alignment of the WhiA C-terminal domain (3hyi, blue), ARC1 (4z8v, green), and Xe4429 (orange). (C) Amino acid sequence alignment of ARC1, Xe4429, and WhiA. Secondary structural and residue number information of ARC1 is shown on top of the alignment. (D) FAM-labelled *XeavrRxo1* promoter DNA and unlabeled *XeavrRxo1* promoter DNA combined with purified proteins, *Xe*AvrRxo1:Xe4429 or Xe4429, were processed via EMSA. The star (*) indicates that the GST-Xe4429 protein bound to the *XeavrRxo1* promoter DNA and caused a mobility shift. Experiments were repeated three times with similar results.

The predicted structure of Xe4429 shares a limited structure homology with the C-terminal domain of DUF199/WhiA from *Thermatoga maritime* (Kaiser *et al*., 2009), although the amino acid sequences of Xe4429, ARC1 (PDB:4z8v), and WhiA (3hyi) were only partially aligned, including the first α-helices from three structures, the third α-helix from Xe4429 and ARC1, and the fourth α-helix from the C-terminal domain of WhiA (Figures 5 B & C). WhiA has been reported as a functional transcription regulator (Kaiser and Stoddard, 2011). A previous report suggests that, in the zeta toxin-antitoxin pairs, the antitoxin can bind to the promoter sequence of the toxin gene to suppress its transcription (Unterholzner *et al*., 2013). Therefore, the antitoxin can also function as a transcriptional repressor. The structural similarity of Xe4429 with WhiA inspired us to test if Xe4429 also functions as a transcription regulator.

### Xe4429 binds to the promoter of XeavrRxo1

To test whether Xe4429 could bind to the promoter region of *XeavrRxo1* and repress the expression of *XeavrRxo1*, we purified either Xe4429 protein alone or the *Xe*AvrRxo1:Xe4429 protein complex and evaluated for binding to a DNA fragment, containing the promoter of *XeavrRxo1* region, using the electrophoretic mobility shift assay (EMSA). The mobility of the DNA fragment containing the *XeavrRxo1* promoter region was delayed when it was incubated with Xe4429 (Figure 5D). The shifting of the DNA band could be inhibited by supplementing unlabeled *XeavrRxo1* promoter DNA as the competitor. In contrast, *Xe*AvrRxo1: Xe4429 protein complex did not alter the mobility of the DNA fragment.

To validate the EMSA results and gain insights into the thermodynamics of the Xe4429-XeAvrRxo1 promoter interaction, we conducted an isothermal titration calorimetry (ITC) assay using an 81 bp DNA fragment encompassing the region immediately adjacent to the start codon of XeAvrRxo1.Analysis of the interaction, considering a stoichiometry of 1 and through data fitting, revealed an estimated dissociation constant (*K*d) of 56 ± 10 µM and a Gibbs free energy constant of –6.0 ± 0.1 kcal/mol. The negative value of the Gibbs free energy constant underscores the thermodynamic favorability of Xe4429 binding to the XeavrRxo1 promoter. These combined findings robustly support the assertion that Xe4429 directly binds to the promoter region of *XeavrRxo1*.

### Xe4429 functions as a transcriptional repressor in X. euvesicatoria cells

Since Xe4429 is able to bind to the promoter of *XeavrRxo1* (Figures 5D and 6), we further investigated whether Xe4429 can also regulate the expression of *XeavrRxo1* in *X. euvesicatoria* cells. The native promoter of *XeavrRxo1* was cloned and fused with the open reading frame of the *luciferase* (*NanoLuc*) gene (England *et al*., 2016). The DNA fragment was subcloned into a pBMTBX-2 (Prior *et al*., 2010) vector that carries either *Xe4429* alone, *XeavrRxo1*-*Xe4429*, or *GFP* genes (Figure 7A). As a control, we also cloned the *luciferase* gene alone into pBMTBX-2 (Figure 7A). The expression constructs were transformed into *XeΔavrRxo1* mutant strain, and grown on the medium supplement with or without arabinose. The bioluminescence signal generated by luciferase was monitored. Expression of Xe4429, but not GFP, repressed the expression of *XeavrRxo1* promoter and suppressed the bioluminescence signal (Figure 7B, panels 2 and 6). Somehow, the bacterial cells carrying the *GFP* construct produced relatively weaker bioluminescence signals. Nevertheless, the induced expression of *GFP* does not affect the bioluminescence signals (panel 6). Interestingly, the expression of *Xe*AvrRxo1:Xe4429 complex also repressed the expression of *Xe*avrRxo1 promoter (Figure 7B, panel 3). This data is inconsistent with the EMSA result shown in Figure 5D, where the *Xe*AvrRxo1:Xe4429 protein complex cannot bind to the promoter of *XeavrRxo1*. It is possible, however, some Xe4429 proteins can be released from the *Xe*AvrRxo1:Xe4429 complex and subsequently regulate the expression of *XeavrRxo1*. We further investigated whether Xe4429 could repress the expression *XeavrRxo1* when *X. euvesicatoria* cells were grown in plant tissues. The *X. euvesicatoria* strains carrying constructs are listed in Figure 7A, and were infiltrated into pepper leaves, supplemented with either arabinose (expression inducer) or glucose (non-inducer). Strong luciferase enzyme activities resulting in a clear bioluminescence signal were observed in pepper leaf inoculated with all *X. euvesicatoria* strains (Figure 7C). Thus, when *X. euvesicatoria* cells are grown in pepper leaf tissues, Xe4429 cannot repress the expression of *XeavrRxo1*.

**Figure 6.**
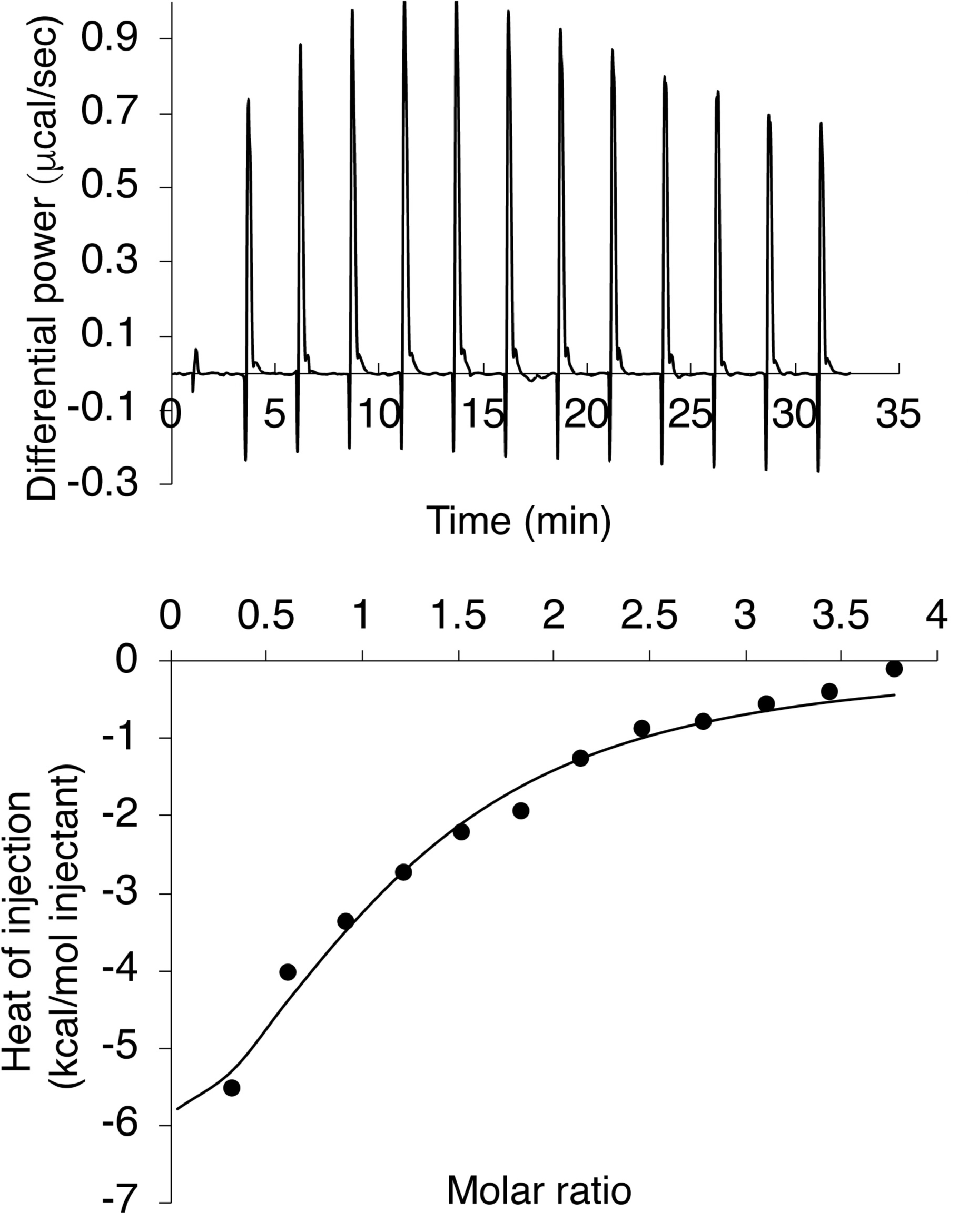
Direct Xe4429 binding to the *XeavrRxo1 promoter*. The upper panel represents the raw titration data, whereas the lower panel shows the integrated, normalized concentration and the corrected heat of dilution data.Fitting was carried outt using a single binding site model and N fixed to 1.

**Figure 7.**
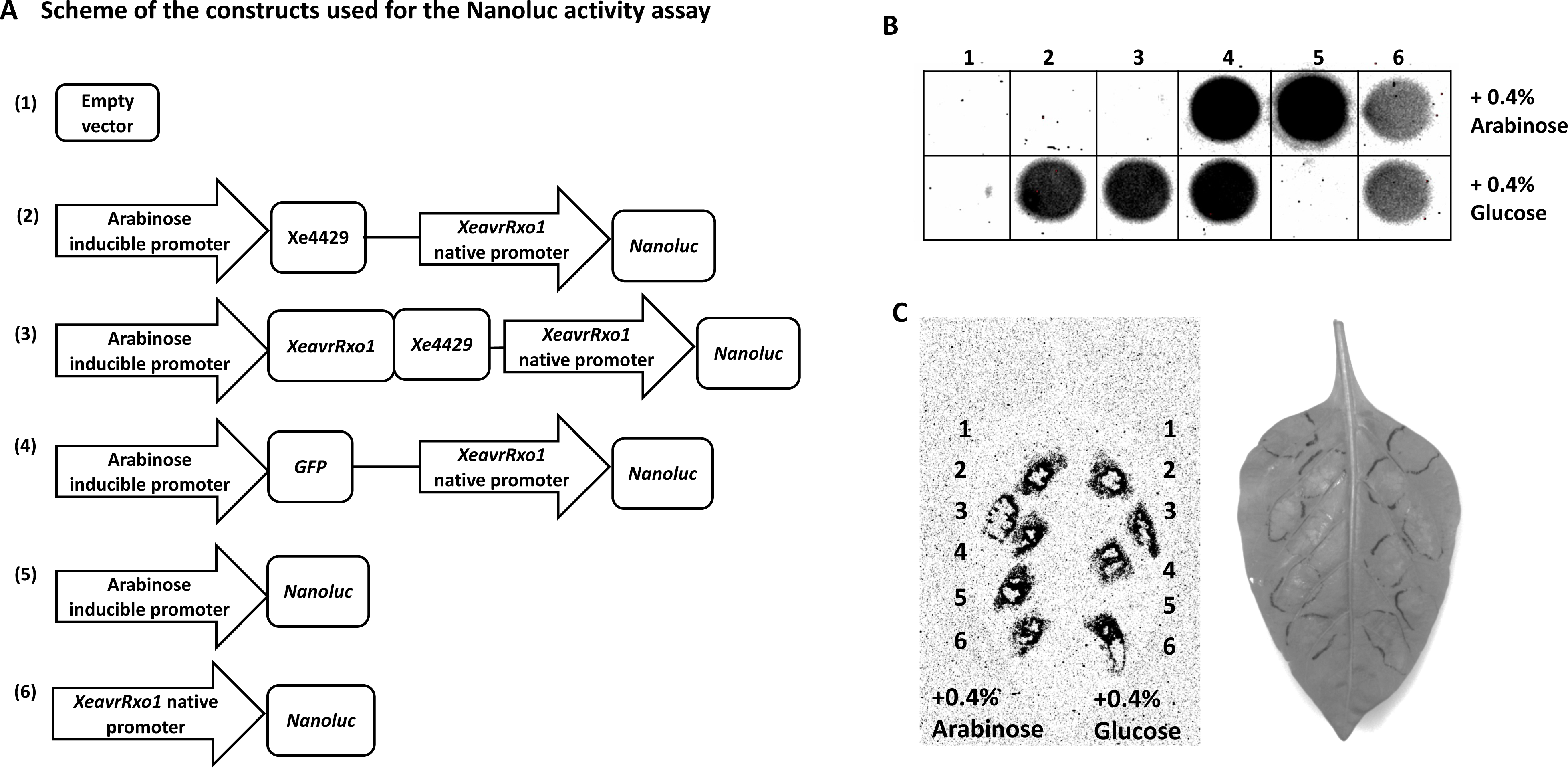
Xe4429 suppresses transcription of *XeavrRxo1* in *X. euvesicatoria* bacterial cells grown in NYG medium but not in plant tissues. (A) Scheme of different constructs used for the luciferase enzymatic activity assay. (B) *X. euvesicatoria* strains carrying the following plasmids: (1) *XeρXeavrRxo1*(pBMTBX-empty vector), (2) *XeρavrRxo1* (pBMTBX-*Xe4429+XeavrRxo1*pro-*NanoLuc*), (3) *XeρXeavrRxo1* (pBMTBX-*XeavrRxo1-Xe4429+XeavrRxo1*pro-*Nanoluc*), (4) *XeρXeavrRxo1* (pVSP61-*XeavrRxo1*pro-*NanoLuc*) (5), *XeρXeavrRxo1* (pBMTBX-*NanoLuc*), and (6) *XeρXeavrRxo1* (pBMTBX-*GFP*-*XeavrRxo1*pro-*NanoLuc*) were grown in NYG medium. The bacterial cells were collected in a 96-well plate and induced with either arabinose or glucose for 6 hours. The luciferase signal was detected using a CCD camera. *XeavrRxo1* promoter-*Nanoluc* is constitutively expressed in *X. euvesicatoria* (panel 4), while the co-expression of Xe4429 or *Xe*AvrRxo1-Xe4429 suppresses its expression (panels 2 and 3). (C) The same set of *X. euvesicatoria* strains were also infiltrated into pepper leaf supplemented with either arabinose or glucose. Leaf samples were collected at 24 h after infiltration. The bioluminescence signal was detected for all inoculated strains. Experiments were conducted at least three times with similar results.

### Expression of Xe4429 enhanced the secretion and translocation of XeAvrRxo1

We previously speculated that *Xoc*AvrRxo1-ORF2 (ARC1) might function as a molecular chaperone that can facilitate the secretion and translocation of *Xoc*AvrRxo1 (Zhao *et al*., 2004). However, we were not able to validate the hypothesis because of the difficulty of generating *XocavrRxo1* mutant in *Xoc*. Here, we investigated the role of Xe4429 during the translocation of *Xe*AvrRxo1. The calmodulin-dependent adenylate cyclase (Cya) reporter assay was conducted in pepper leaf tissues (Figure 8). The non-toxic *XeavrRxo1* (D222T) gene was fused to *Cya* gene and integrated into the genome of *XeΔavrRxo1*. The derived strain was designated as *Xe*-D222T: Cya. The *Xe4429* gene, driven by an arabinose inducible promoter, was transformed into *Xe*-D222: Cya. Xcv85-10 strain carries an *avrBs2*: *Cya* reporter gene was used as a positive control in the Cya assay (Zhao *et al*., 2011). Although *Xe*AvrRxo1(D222T): Cya alone could be translocated into plant cells with increased Cya activities, co-expression with *Xe4429* could significantly increase Cya signals of *Xe*AvrRxo1(D222T): Cya (Figure 8). Thus, Xe4429 can function as a translocation facilitator to enhance the translocation of *Xe*AvrRxo1 into plant cells.

**Figure 8.**
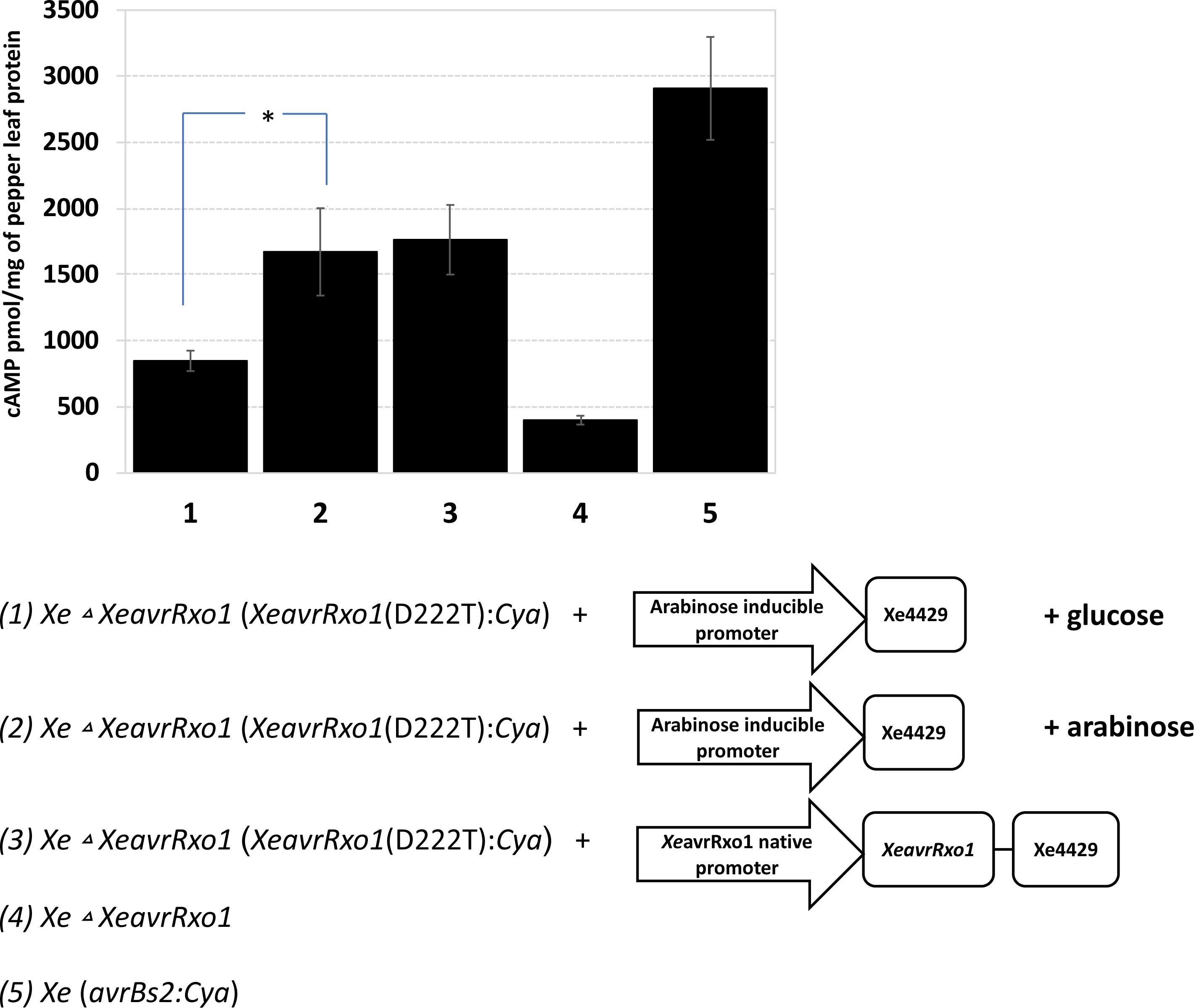
Calmodulin-dependent adenylate cyclase (Cya) reporter assay shows that Xe4429 facilitates the translocation of *Xe*AvrRxo1 into pepper plant cells. *XeρavrRxo1* strains carrying the indicated Cya reporter (were used for Cya assay. (1). *Xe*-D222T:Cya + pBMTBX-*Xe4429* (0.4 % glucose), (2). *Xe*-D222T:Cya + pBMTBX-*Xe4429* (0.4 % arabinose), (3). *Xe*-D222T:Cya + pBMTBX-*XeAvrRxo1*-*Xe4429* (0.4 % arabinose), (4) *Xe*Δ*XeavrRxo1*, (5) *Xe*(*avrBs2:Cya*). Cya activity was measured and indicated by cAMP production (pmol/mg of leaf protein) in pepper leaves infiltrated with the *X. euvesicatoria* strains (4 x 10^8^ CFU ml^-1^). Leaf samples were collected 8 hours after infiltration. Values are the average of three replicates, and the error bars represent standard errors.

## Discussion

### XeAvrRxo1 is a functional orthologue of XocAvrRox1 with both virulence and avirulence functions

We characterized a T3E *Xe*AvrRxo1 from *X. euvesicatoria* strain Xcv85-10. *Xe*AvrRxo1 is a functional orthologue of *Xoc*AvrRxo1, which can trigger an Rxo1-mediated hypersensitive response in *N. benthamiana* and pepper plants (Figures 1 and 2). Interestingly, another *avrRxo1* homolog gene, *Aave_3602*, cloned from the watermelon pathogen *Acidovorax citrulli,* cannot trigger the Rxo1-mediated hypersensitive response (Figure S3A). Aave_3602 shares 55% of its identities with XocAvrRxo1 and 59% with *Xe*AvrRxo1. The ATP-binding motif and the NAD substrate binding site are conserved in Aave_3602. A few amino acids that are conserved in *Xoc*AvrRxo1 and *Xe*AvrRxo1, but not in Aave_3602, could be essential for the interaction of Rxo1 and AvrRxo1 (Figure S3B).

We previously demonstrated that the maize *Rxo1* gene can be functionally transferred into rice and condition disease resistance to bacterial leaf streak disease caused by *X. o. oryzicola* (Zhao *et al*., 2005). It demonstrates the feasibility of transferring a nonhost *R* gene between cereals and providing a valuable tool for controlling bacterial leaf streak disease. However, the molecular mechanism of Rox1/AvrRxo1-mediated disease resistance is still elusive. It is also unclear if we can transfer *Rxo1* to the far-related plant species. Therefore, it is appealing to develop stable transgenic tobacco or pepper plants expressing the maize *Rxo1* gene and test if *Rxo1* also conditions disease resistance to *X. euvesicatoria* in dicot plant species, which diverged from monocot species about 140–200 million years ago (Chaw *et al*., 2004; Wolfe *et al*., 1989).

Gram-negative bacterial strains can deliver more than 50 T3Es inside plant cells (Jiménez-Guerrero *et al*., 2020). Most of T3Es have virulence functions that can enhance bacterial proliferation in susceptible host plants. However, the virulence functions of T3Es are redundant, and deletion of a single *T3E* usually does not compromise bacterial virulence (Büttner, 2016). Only a small portion of known T3Es have been confirmed to have significant virulence functions (Büttner, 2016). This subset of T3Es may target defense-related pathways that do not overlap with other co-existing T3Es. Therefore, characterizing the virulence targets of T3Es with significant virulence functions may allow us to identify unique plant immunity components (Block *et al*., 2008; Dillon *et al*., 2019). This study demonstrated that *Xe*AvrRxo1 belongs to the T3Es with strong, non-redundant virulence functions that can promote bacterial growth in pepper and *N. benthamiana* (Figure 3). A previous study suggests that *Xoc*AvrRxo1 also has a significant virulent function that can increase the proliferation of *Xoc* in infected rice plants (Han *et al*., 2015). Therefore, AvrRxo1 and its orthologue may target immunity-related proteins or signaling pathways conserved in both monocot and dicot plant species. It will be intriguing to characterize the virulence mechanism of AvrRxo1 further. We previously failed to develop an *avrRxo1*-deletion mutant in *Xoc*, which limits our ability to further characterize the biological and biochemical functions of AvrRxo1 in the rice/*Xoc* pathosystem (Zhao *et al*., 2004). In this study, we successfully deleted both *Xe*AvrRxo1 and its putative chaperone Xe4429 in the genome of *X. euvesicatoria* strain Xcv85-10 (Figure 3). Validation of the virulence and avirulence functions of *Xe*AvrRxo1 in pepper and *N. benthamiana*/ *X. euvesicatoria* pathosystem enables us to further explore the biological and biochemical functions of AvrRxo1 and Xe4429.

### Conserved gene structure of avrRxo1:ARC1

In the *Xoc* genome, *avrRxo1* is adjacent to another gene, *ARC1* (previously named AvrRxo1-ORF2), which encodes a molecular chaperone of AvrRxo1 (Han *et al*., 2015). This study shows that *XeavrRxo1* is located near an ARC1 homolog, referred to as Xe4429. We previously demonstrated that overexpression of AvrRxo1 suppresses bacterial growth (Triplett *et al*., 2016). The cytotoxicity of AvrRxo1 can be suppressed by co-expression with ARC1 (Han *et al*., 2015; Triplett *et al*., 2016). Like *Xoc*AvrRxo1, the inducible expression of *Xe*AvrRxo1 in *E. coli* and *X. euvesicatoria* cells is toxic, and the co-expression with Xe4429 can suppress the toxicity. However, the cell growth of *X. euvesicatoria* and *E. coli* is still partially impacted (Figures 4A and B). Notably, the expression of *XeavrRxo1* and *Xe4429* under their native promoter did not affect the cell proliferation of *X. euvesicatoria*. (Figure 4C). We hypothesize that the expression of *XeAvrRxo1-Xe4429* genes under an arabinose-inducible promoter could lead to an association-dissociation dynamic within the XeAvrRxo1:Xe4429 complex, potentially resulting in the release of free XeAvrRxo1, which may exhibit toxicity towards bacterial cells. However, when its native promoter expresses the *XeavrRxo1*-*Xe4429* genes, some free Xe4429 could bind to the promoter of *XeavRxo1* and reduce its transcription/translation (Figure 5D). This also explains why the *Xe*AvrRxo1: Xe4429 protein complex can suppress the expression of *XeavrRxo1* promoter-luciferase (Figure 7B). This translation-responsive model is also reported in the type II toxin-antitoxin, e.g., the ε/ζ systems (Chan *et al*., 2016; Mutschler and Meinhart, 2011). However, the gene structure of *XeavrRxo1*-*Xe4429* is different from the ε/ζ system, where the antitoxin is located upstream of the toxin gene (Mutschler and Meinhart, 2011). Nevertheless, we revealed an autoregulation mechanism of *Xe*AvrRxo1: Xe4429. In the *X. euvesicatoria* genome, *XeavrRxo1-Xe4429* genes are organized as an operon that is controlled by the *XeAvrRxo1* promoter. The expressed *Xe*AvrRxo1:Xe4429 proteins can form a complex that is non-toxic to *X. euvesicatoria* cells. Some free Xe4429 proteins can bind to the promoter of *XeAvrRxo1* and repress the transcription of *XeavrRxo1*. This translation-responsive phenomenon was not observed in the *Xoc* system because of the lack of an *avrRxo1*-deletion mutant. This is also the first time demonstrating that a *T3E* can be autoregulated in a translation-responsive model. A few *T3E* genes in the genomes of phytopathogens are adjacent to other small open reading frames, which may function as chaperone proteins (Lohou *et al*., 2013). It will be interesting to investigate whether those *T3E* genes are regulated by their cognate chaperones in a manner similar to this study.

### One gene with three functions: Xe4429 encodes as an antitoxin, a transcriptional/translational repressor, and a facilitator of T3E-translocation

As illustrated in Figure 9, we have identified that the Xe4429 protein serves a triple role, intricately intertwining to finely regulate the expression, toxicity, and translocation of XeAvrRxo11. In addition to the antitoxin function, we identified Xe4429 also functions as a transcription/translation repressor in *X. euvesicatoria* cells (Figures 5D, 6, and 7B). We utilized two distinct methodologies to assess the molecular interaction between Xe4429 and the XeAvrRxo1 promoter DNA fragments. Although EMSA is a widely adopted technique offering a rapid and qualitative assessment of DNA-protein binding, it is limited for quantitarive purposes. Therefore, using ITC, we show that Xe4429 binds to an 81 bp fragment of the XeAvrRxo1 promoter with a micromolar affinity. This protein-DNA interaction may potentially lead to the complete inhibition of XeAvrRxo1transcription and/or translation, consistent with the findings in Figure 7, where the co-expression of Xe4429 suppresses the expression of the Nanoluc gene driven by the *XeAvrRxo1* promoter. However, during the infection of *X. euvesicatoria* to pepper leaves, the Xe4429-mediated repression is released (Figure 7C). It will be interesting to further investigate the bio and abio-factors that can suppress the Xe4429-mediated repression in plant tissues. We also demonstrate that Xe4429 can enhance the secretion and translocation of *Xe*AvrRxo1(D222T) (Figure 8). Our previous work suggests that ARC1 (Xoc-AvrRxo1-ORF2) is not required for the delivery of *Xoc*AvrRxo1 (Zhao *et al*., 2004). Consistent with previous studies, *Xe*AvrRxo1 (D222T) is delivered into plant cells even without the assistance of Xe4429. However, the co-expression of Xe4429 can significantly enhance the translocation rates of *Xe*AvrRxo1(D222T), resulting in increased Cya activities (Figure 8). Therefore, Xe4429 may function as a translocation facilitator during the *X. euvesicatoria* pathogenesis. It is still unclear whether Xe4429 can also enhance the translocation of other T3Es in *X. euvesicatoria*, which may be worthy of further investigation in the future.

**Figure 9.**
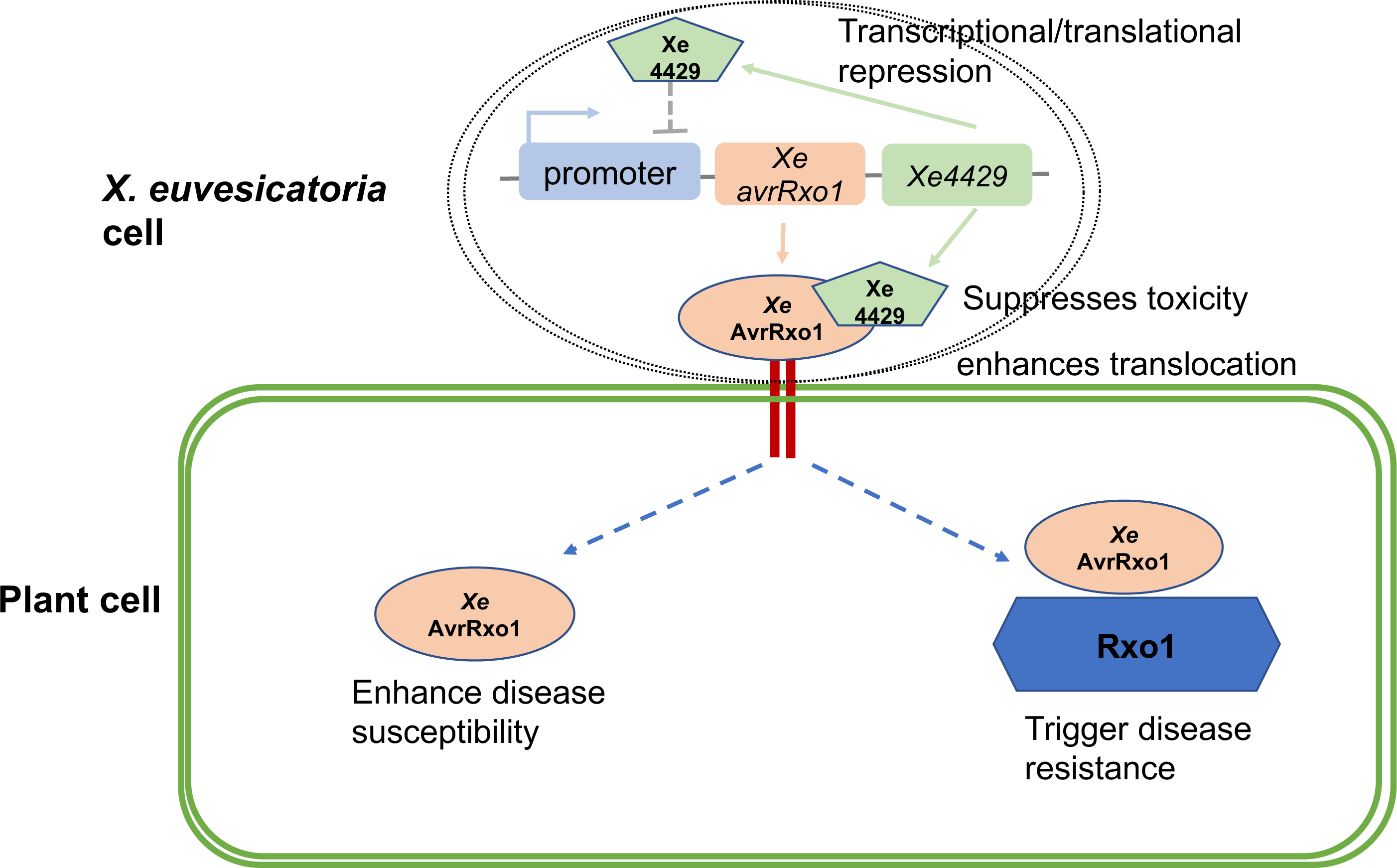
The working model of Xe4429 with three biological and biochemical functions. Inside *X. euvesicatoria* bacterial cells, Xe4429 directly interacts with *Xe*AvrRxo1 and functions as an antitoxin. Xe4429 also binds to the promoter of *XeavrRxo1* and suppresses its transcription. The Xe4429-mediated repression is released upon the infection of *X. euvesicatoria* to plant cells. In addition, during plant cell invasion, Xe4429 promotes the translocation of *Xe*AvrRxo1, therefore enhancing the virulence of *X. euvesicatoria*. Maize Rxo1 can recognize *Xe*AvrRxo1 to trigger defense responses.

## Acknowledgments

This work was supported by the US National Science Foundation (IOS-0845283) to BZ, the Binational Agricultural Research and Development Fund (IS-5023-17C) to BZ, and the Virginia Agricultural Experiment Station (VA-160144) to BZ.

## Author contributions

ZW, CZ, and BZ designed the research; ZW, CZ, TGR, QL, KW, JM, CT, SW, YT, QH, and FS, and JL performed the experiments; ZW, CZ, TGR, DGSC, JL, and BZ analyzed the data; ZW, DGSC, TGR, and BZ wrote the paper with input from all the authors.

## Figure legends

**Figure S1.**
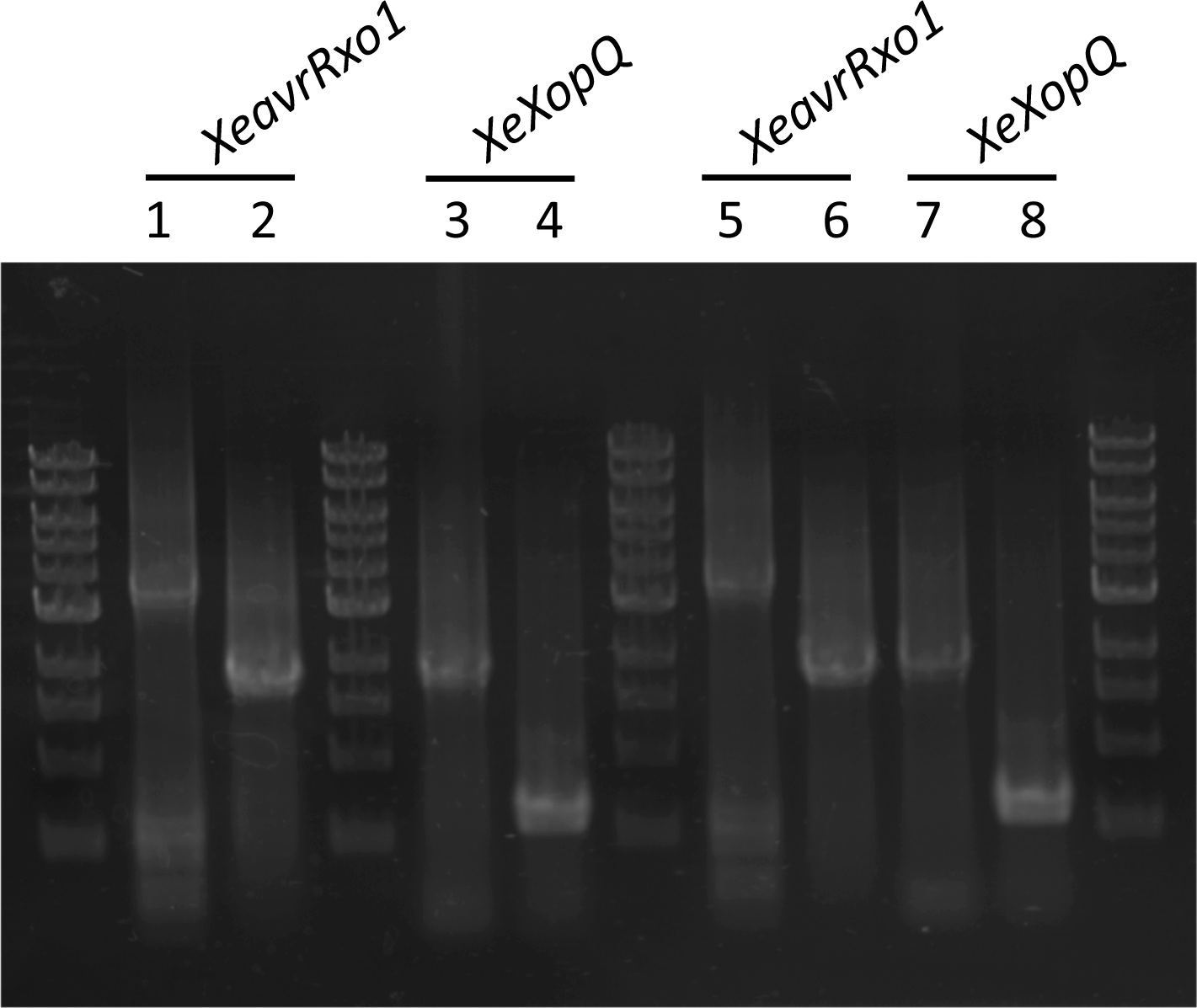
The deletion of *XopQ* and *XeavrRxo1-Xe4429* was validated by PCR amplification with the primers flanking the targeted genes. Lane 1, 3, 5, 7: Wild type *X. euvesicatoria strain Xcv85-10,* lane 2: *Xe△XeavrRxo1*, lane 4: *Xe△XopQ*, lane 6 and 8: *Xe△XeavrRxo1△XopQ*. *XeavrRxo1* flanking DNA was amplified with primers Xe4428pro For and Xcv4430Rev. *XopQ* flanking DNA was amplified with primers Xcv4438 checkfor and Xcv4438 checkrev. M: 1Kb DNA maker.

**Figure S2.**
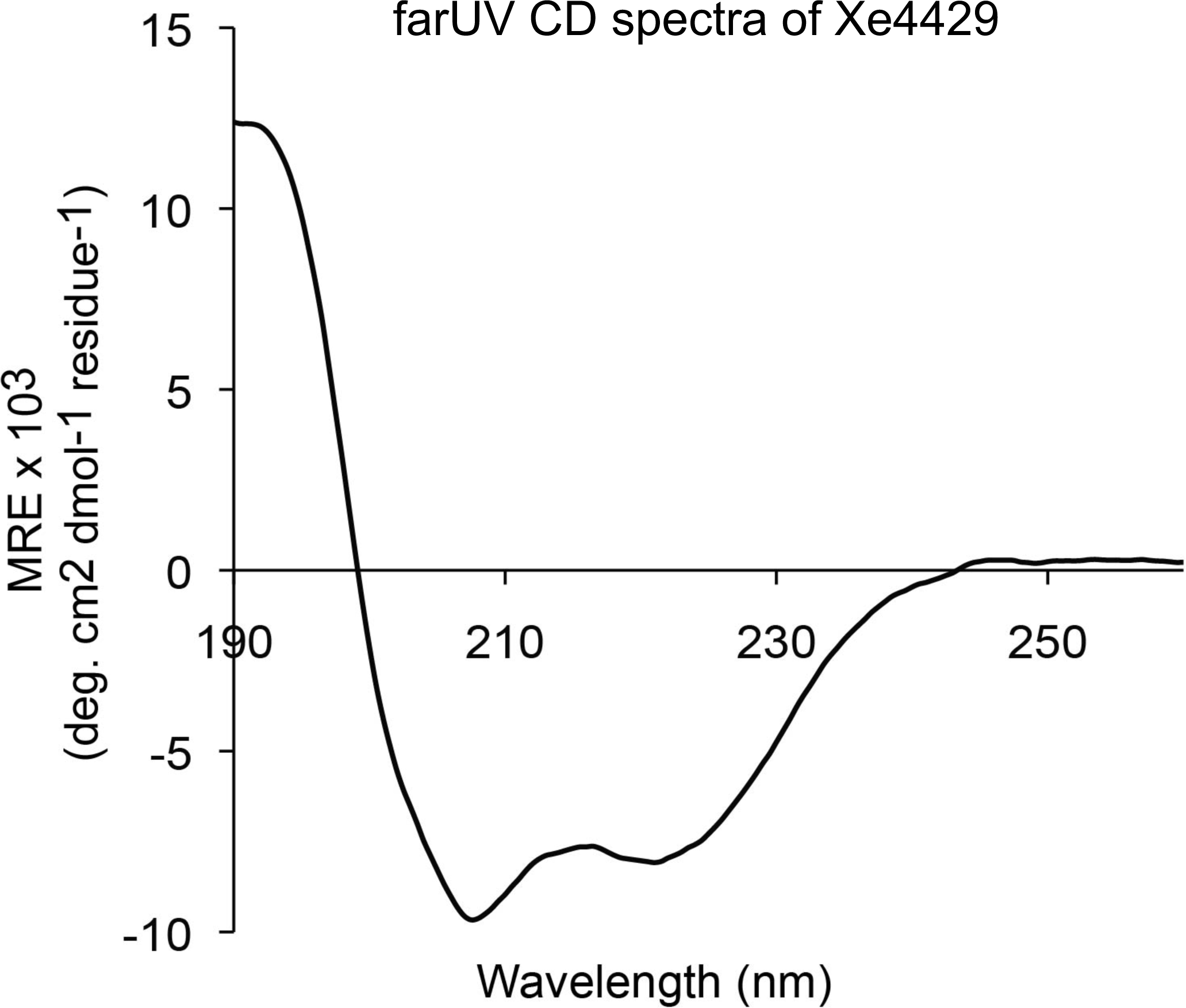
Far-UV circular dichroism spectrum of Xe4429. The spectrum displays two well-resolved minima at 208 and 221 nm, indicating that Xe4429 is largely α-helical.

**Figure S3.**
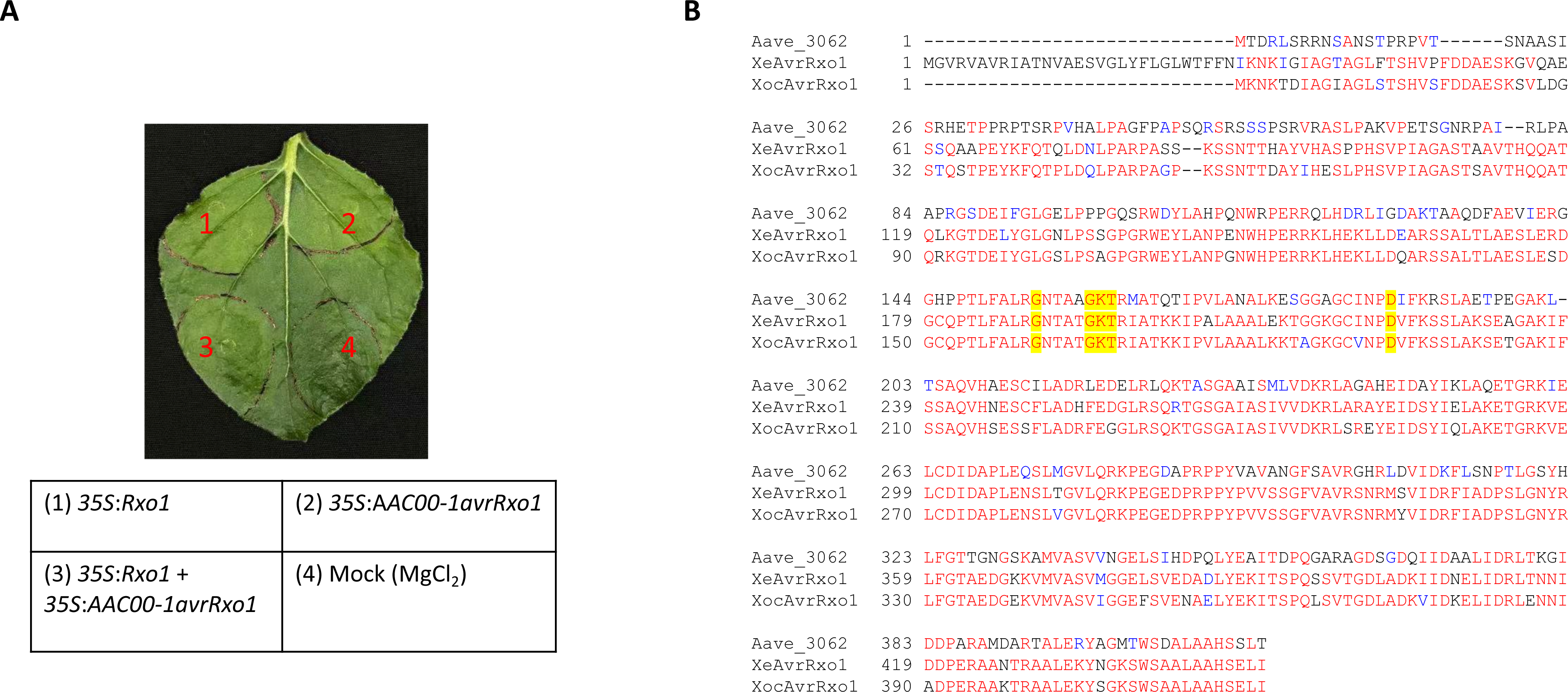
Transient co-expression of Aave_3602 with Rxo1 does not trigger cell death in *N. benthamiana*. The open reading frame of *Aave_3602* was amplified from the genome DNA of *Acidovorax citrulli* strain AAC00-1 with primers Aave_3062 BamFor1 and Aave_3062 SalRev1. The PCR product was cloned into pENTR/D-TOPO, and then subcloned into pEarleygate101. Agrobacterium strain GV2260 carrying pEarleygate101-*Aave_3602* was used for transient assay on *N. benthamiana* plant leaves. 35S:*Aave_3602* alone or co-expression with 35S:*Rxo1* failed to trigger cell death phenotype. Picture was taken at two days post-inoculation.

---

**Table S1.**
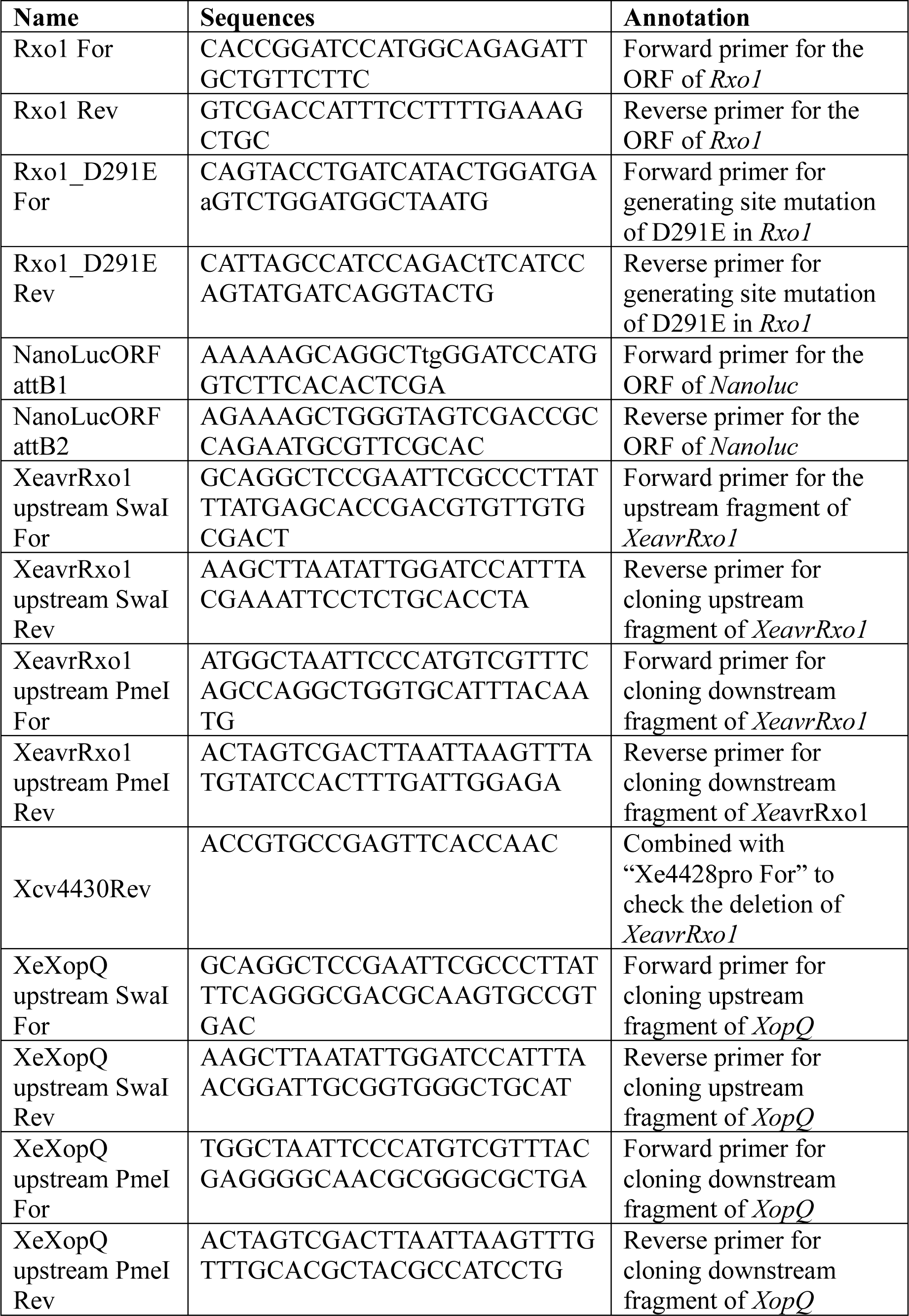
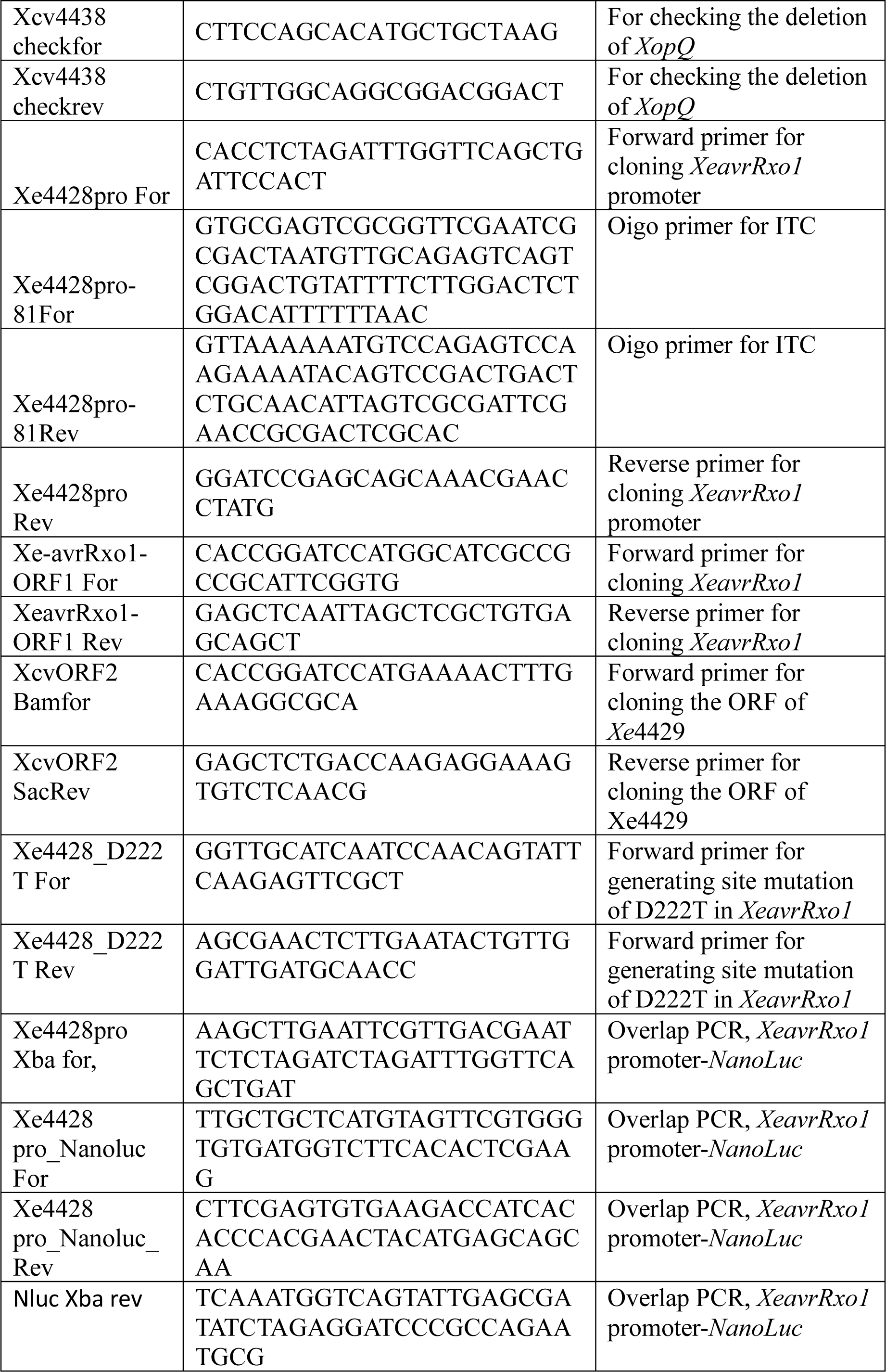
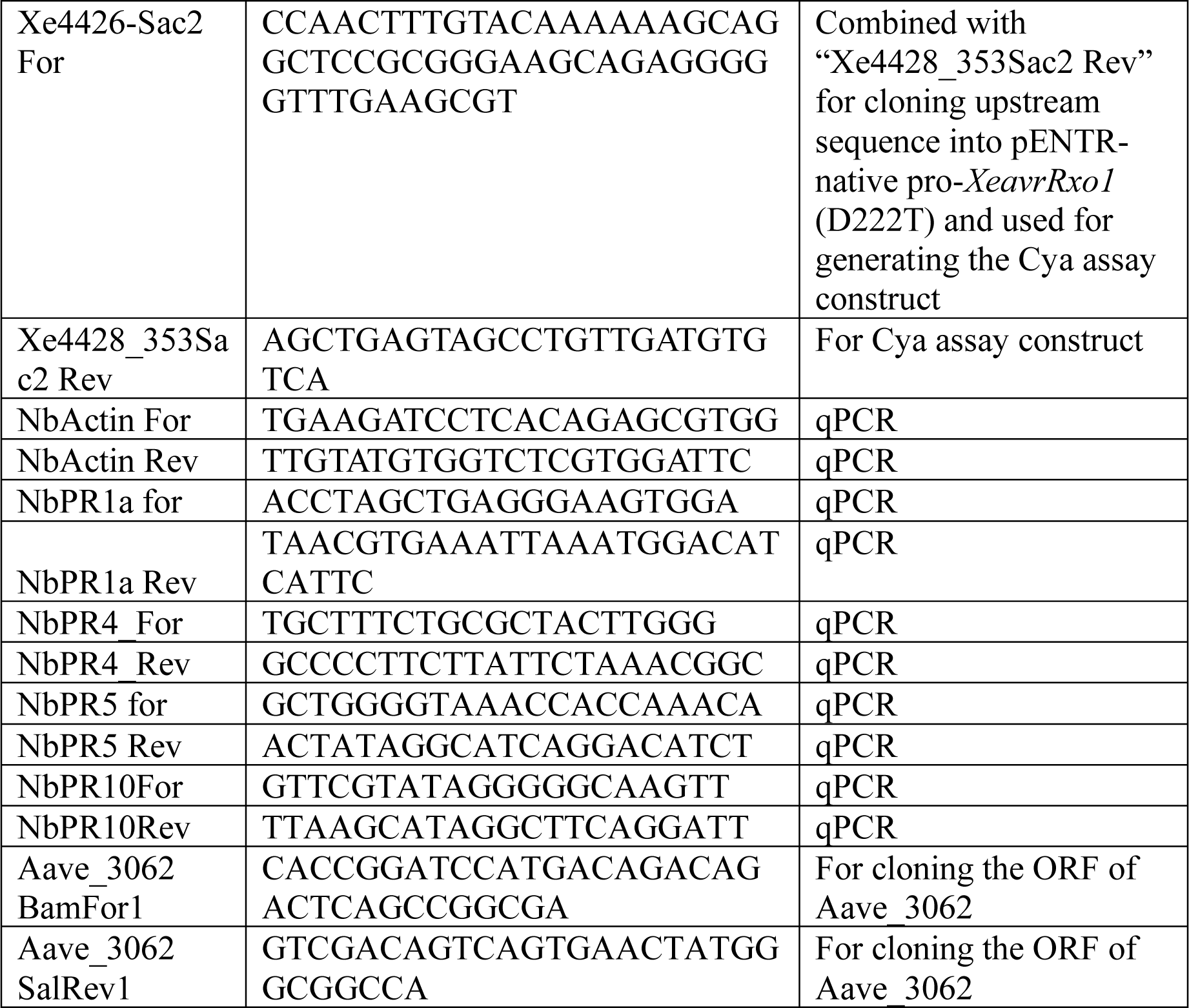
Primers utilized in this study.

## References

1. Alfano JR, Charkowski AO, Deng WL, Badel JL, Petnicki-Ocwieja T, van Dijk K, Collmer A. 2000. The *Pseudomonas syringae* Hrp pathogenicity island has a tripartite mosaic structure composed of a cluster of type III secretion genes bounded by exchangeable effector and conserved effector loci that contribute to parasitic fitness and pathogenicity in plants. Proceedings of the National Academy of Sciences of the United States of America 97, 4856–4861.

2. Arnold DL, Jackson RW, Fillingham AJ, Goss SC, Taylor JD, Mansfield JW, Vivian A. 2001. Highly conserved sequences flank avirulence genes: isolation of novel avirulence genes from *Pseudomonas syringae* pv. *pisi*. Microbiology (Reading) 147, 1171–1182.

3. Block A, Li G, Fu ZQ, Alfano JR. 2008. Phytopathogen type III effector weaponry and their plant targets. Current opinion in plant biology 11, 396–403.

4. Büttner D. 2016. Behind the lines–actions of bacterial type III effector proteins in plant cells. FEMS Microbiology Reviews 40, 894–937.

5. Casper-Lindley C, Dahlbeck D, Clark ET, Staskawicz BJ. 2002. Direct biochemical evidence for type III secretion-dependent translocation of the AvrBs2 effector protein into plant cells. Proceedings of the National Academy of Sciences of the United States of America 99, 8336–8341.

6. Chan WT, Espinosa M, Yeo CC. 2016. Keeping the wolves at bay: antitoxins of prokaryotic type II toxin-antitoxin systems. Frontiers in Molecular Biosciences 3.

7. Chaw S-M, Chang C-C, Chen H-L, Li W-H. 2004. Dating the monocot–dicot divergence and the origin of core eudicots using whole chloroplast genomes. Journal of Molecular Evolution 58, 424–441.

8. Dillon MM, Almeida RND, Laflamme B, Martel A, Weir BS, Desveaux D, Guttman DS. 2019. Molecular evolution of *Pseudomonas syringae* type III secreted effector proteins. Frontiers in Plant Science 10.

9. Earley KW, Haag JR, Pontes O, Opper K, Juehne T, Song K, Pikaard CS. 2006. Gateway-compatible vectors for plant functional genomics and proteomics. Plant Journal 45, 616–629.

10. England CG, Ehlerding EB, Cai W. 2016. NanoLuc: A small luciferase is brightening up the field of bioluminescence. Bioconjugate chemistry 27, 1175–1187.

11. Gille C, Frömmel C. 2001. STRAP: editor for STRuctural Alignments of proteins. Bioinformatics 17, 377–378.

12. Han Q, Zhou C, Wu S, et al. 2015. Crystal structure of *Xanthomonas* AvrRxo1-ORF1, a type III effector with a polynucleotide kinase domain, and its interactor AvrRxo1-ORF2. Structure 23, 1900–1909.

13. Jiménez-Guerrero I, Pérez-Montaño F, Da Silva GM, et al. 2020. Show me your secret(ed) weapons: a multifaceted approach reveals a wide arsenal of type III-secreted effectors in the cucurbit pathogenic bacterium Acidovorax citrulli and novel effectors in the Acidovorax genus. Molecular Plant Pathology 21, 17–37.

14. Jones JB, Lacy GH, Bouzar H, Stall RE, Schaad NW. 2004. Reclassification of the xanthomonads associated with bacterial spot disease of tomato and pepper.

15. Kaiser BK, Clifton MC, Shen BW, Stoddard BL. 2009. The structure of a bacterial DUF199/WhiA protein: domestication of an invasive endonuclease. Structure 17, 1368–1376.

16. Kaiser BK, Stoddard BL. 2011. DNA recognition and transcriptional regulation by the WhiA sporulation factor. Scientific Reports 1, 156.

17. Levy A, Salas Gonzalez I, Mittelviefhaus M, et al. 2018. Genomic features of bacterial adaptation to plants. Nature Genetics 50, 138–150.

18. Liu Y, Miao J, Traore S, Kong D, Liu Y, Zhang X, Nimchuk ZL, Liu Z, Zhao B. 2016. SacB-SacR gene cassette as the negative selection marker to suppress *Agrobacterium* Overgrowth in Agrobacterium-mediated plant transformation. Frontiers in Molecular Biosciences 3, 70–70.

19. Liu Y, Wang K, Cheng Q, et al. 2020. Cysteine protease RD21A regulated by E3 ligase SINAT4 is required for drought-induced resistance to *Pseudomonas syringae* in Arabidopsis. Journal of Experimental Botany 71, 5562–5576.

20. Liu Y, Zhang X, Tran H, Shan L, Kim J, Childs K, Ervin EH, Frazier T, Zhao B. 2015. Assessment of drought tolerance of 49 switchgrass (*Panicum virgatum*) genotypes using physiological and morphological parameters. Biotechnol Biofuels 8, 152.

21. Lohou D, Lonjon F, Genin S, Vailleau F. 2013. Type III chaperones & Co in bacterial plant pathogens: a set of specialized bodyguards mediating effector delivery. Frontiers in Plant Science 4.

22. Mutschler H, Meinhart A. 2011. ε/ζ systems: their role in resistance, virulence, and their potential for antibiotic development. Journal of molecular medicine (Berlin, Germany) 89, 1183–1194.

23. Popov G, Fraiture M, Brunner F, Sessa G. 2016. Multiple *Xanthomonas euvesicatoria* type III effectors inhibit flg22-triggered immunity. Molecular Plant-Microbe Interactions 29, 651–660.

24. Potnis N, Timilsina S, Strayer A, Shantharaj D, Barak JD, Paret ML, Vallad GE, Jones JB. 2015. Bacterial spot of tomato and pepper: diverse *Xanthomonas* species with a wide variety of virulence factors posing a worldwide challenge. Molecular Plant Pathology 16, 907–920.

25. Prior JE, Lynch MD, Gill RT. 2010. Broad-host-range vectors for protein expression across gram negative hosts. Biotechnol Bioeng 106, 326–332.

26. Roden JA, Belt B, Ross JB, Tachibana T, Vargas J, Mudgett MB. 2004. A genetic screen to isolate type III effectors translocated into pepper cells during *Xanthomonas infection*. Proceedings of the National Academy of Sciences of the United States of America 101, 16624–16629.

27. Schwartz AR, Potnis N, Timilsina S, et al. 2015. Phylogenomics of *Xanthomonas* field strains infecting pepper and tomato reveals diversity in effector repertoires and identifies determinants of host specificity. Frontiers in Microbiology 6.

28. Shidore T, Broeckling CD, Kirkwood JS, Long JJ, Miao J, Zhao B, Leach JE, Triplett LR. 2017. The effector AvrRxo1 phosphorylates NAD in planta. PLOS Pathogens 13, e1006442.

29. Stall R, Beaulieu C, Egel D, Hodge N, Leite R, Minsavage G, Bouzar H, Jones J, Alvarez A, Benedict A. 1994. Two genetically diverse groups of strains are included in *Xanthomonas campestris* pv. *vesicatoria*. International Journal of Systematic and Evolutionary Microbiology 44, 47–53.

30. Stall RE, Jones JB, Minsavage GV. 2009. Durability of resistance in tomato and pepper to xanthomonads causing bacterial spot. Annual review of phytopathology 47, 265–284.

31. Teper D, Burstein D, Salomon D, Gershovitz M, Pupko T, Sessa G. 2016. Identification of novel *Xanthomonas euvesicatoria* type III effector proteins by a machine-learning approach. Molecular Plant Pathology 17, 398–411.

32. Thieme F, Koebnik R, Bekel T, et al. 2005. Insights into genome plasticity and pathogenicity of the plant pathogenic bacterium Xanthomonas campestris pv. vesicatoria revealed by the complete genome sequence. Journal of bacteriology 187, 7254–7266.

33. Traore SM, Eckshtain-Levi N, Miao J, et al. 2019. *Nicotiana* species as surrogate host for studying the pathogenicity of *Acidovorax citrulli*, the causal agent of bacterial fruit blotch of cucurbits. Molecular Plant Pathology 20, 800–814.

34. Triplett LR, Shidore T, Long J, et al. 2016. AvrRxo1 Is a Bifunctional Type III Secreted Effector and Toxin-Antitoxin System Component with Homologs in Diverse Environmental Contexts. PLoS One 11, e0158856.

35. Unterholzner SJ, Poppenberger B, Rozhon W. 2013. Toxin-antitoxin systems: Biology, identification, and application. Mobile Genet Elements 3, e26219.

36. Vallejos CE, Jones V, Stall RE, Jones JB, Minsavage GV, Schultz DC, Rodrigues R, Olsen LE, Mazourek M. 2010. Characterization of two recessive genes controlling resistance to all races of bacterial spot in peppers. Theoretical and applied genetics 121, 37–46.

37. Van Eck L, Davidson RM, Wu S, Zhao BY, Botha AM, Leach JE, Lapitan NL. 2014. The transcriptional network of WRKY53 in cereals links oxidative responses to biotic and abiotic stress inputs. Functional & Integrative Genomics 14, 351–362.

38. Vauterin L, Hoste B, Kersters K, Swings J. 1995. Reclassification of xanthomonas. International Journal of Systematic and Evolutionary Microbiology 45, 472–489.

39. Williams CJ, Headd JJ, Moriarty NW, et al. 2018. MolProbity: More and better reference data for improved all-atom structure validation. Protein Science 27, 293–315.

40. Wolfe KH, Gouy M, Yang YW, Sharp PM, Li WH. 1989. Date of the monocot-dicot divergence estimated from chloroplast DNA sequence data. Proceedings of the National Academy of Sciences of the United States of America 86, 6201–6205.

41. Yang JT, Wu CS, Martinez HM. 1986. Calculation of protein conformation from circular dichroism. Methods in Enzymology 130, 208–269.

42. Zhao B, Ardales EY, Raymundo A, Bai J, Trick HN, Leach JE, Hulbert SH. 2004. The *avrRxo1* gene from the rice pathogen *Xanthomonas oryzae* pv. *oryzicola* confers a nonhost defense reaction on maize with resistance gene *Rxo1*. Molecular Plant-Microbe Interactions 17, 771–779.

43. Zhao B, Dahlbeck D, Krasileva KV, Fong RW, Staskawicz BJ. 2011. Computational and biochemical analysis of the *Xanthomonas* effector AvrBs2 and its role in the modulation of *Xanthomonas* type three effector delivery. PLOS Pathogens 7, e1002408.

44. Zhao B, Lin X, Poland J, Trick H, Leach J, Hulbert S. 2005. A maize resistance gene functions against bacterial streak disease in rice. Proceedings of the National Academy of Sciences of the United States of America 102, 15383–15388.

